# Universal correction of enzymatic sequence bias reveals molecular signatures of protein/DNA interactions

**DOI:** 10.1101/104364

**Authors:** André L. Martins, Ninad M. Walavalkar, Warren D. Anderson, Chongzhi Zang, Michael J. Guertin

## Abstract

Coupling molecular biology to high throughput sequencing has revolutionized the study of biology. Molecular genomics techniques are continually refined to provide higher resolution mapping of nucleic acid interactions and structure. Sequence preferences of enzymes can interfere with the accurate interpretation of these data. We developed *seqOutBias* to characterize enzymatic sequence bias from experimental data and scale individual sequence reads to correct intrinsic enzymatic sequence biases. *SeqOutBias* efficiently corrects DNase-seq, TACh-seq, ATAC-seq, MNase-seq, and PRO-seq data. We show that *seqOutBias* correction facilitates identification of true molecular signatures resulting from transcription factors and RNA polymerase interacting with DNA.

## Introduction

The field of molecular genomics emerged as classical molecular biology techniques were coupled to high throughput sequencing technology to provide unprecedented genome-wide measurements of molecular features. Molecular genomics assays, such as DNase-seq (1, 2), ChIP-exo (3), and PRO-seq (4, 5), are converging on single-nucleotide resolution measurements. The enzymes that are routinely used in molecular biology and cloning have inherent and often uncharacterized sequence preferences. These preferences manifest more prominently as the resolution of genomic assays increases. Therefore, we developed *seqOutBias* (<u>https://github.com/guertinlab/seqOutBias</u>) to characterize and correct enzymatic biases that can obscure proper interpretation of molecular genomics data.

Enzymatic hypersensitivity assays, such as DNase-seq (1, 2), TACh-seq (6), and ATAC-seq (7), have the potential to measure transcription factor (TF) binding sites genome-wide in a single experiment. These assays strictly measure enzymatic (DNase, Tn5 transposase, Benzonase, or Cyanase) accessibility to DNA and not a specific biological event, making data challenging to deconvolve. Standard algorithms scan for footprints, which are depletions of signal in larger regions of hypersensitivity (8–12). Many transcription factors, however, do not exhibit composite footprints if enzymatic cut frequency is averaged at all ChIP-seq validated binding sites with strong consensus motifs (10–13). Moreover, the inability to detect a footprint at any individual TF binding site results in high false negative rates for footprinting algorithms (14). Accurate footprinting is also confounded by the artifactual molecular signatures that result from enzymatic sequence preference (10–12). DNase footprinting algorithms can incorporate DNase cut preference data to abrogate this bias (12, 15). However, no existing tools specialize in correcting intrinsic sequence bias for a diverse set of enzymes and experimental methodologies.

We find that correcting for enzymatic sequence bias highlights true molecular signatures that result from TF/DNA interactions. Despite the limitations of enzymatic hypersensitivity footprinting and sequence bias signatures, hypersensitive regions reveal a near-comprehensive set of functional regulatory regions in the genome (16). Therefore, we present *seqOutBias*, which calculates sequence bias from an aligned BAM file and corrects individual reads accordingly. While this software does not directly infer transcription factor binding, correction of sequence bias provides a more accurate measurement of three key features of enzymatic hypersensitivity data: 1) raw peak height; 2) footprint depth; and 3) true molecular signatures. These measurements, taken together with DNA sequence, can be used to develop algorithms that infer TF binding genome-wide. Moreover, footprint depth and the presence of true molecular signatures are unique to each TF and these features should be characterized for each TF using corrected data in order to optimize TF-binding inference algorithms.

Enzymatic sequence biases are most well-characterized for DNase-seq experiments (10–12), but nearly all molecular genomics experiments employ enzymatic treatments and these enzymes also have intrinsic biases. Herein, we show that DNase, Cyanase, Benzonase, MNase, Tn5 transposase, and T4 RNA ligase all exhibit sequence preferences that are effectively corrected with *seqOutBias*. We also characterize enzymatic bias that results from T4 DNA Polymerase, T4 Polynucleotide Kinase, and Klenow Fragment (3’→5’ exo-) treatment of DNA in preparation of high throughput sequencing libraries. Lastly, we show that correction of enzymatic sequence bias highlights true molecular signatures, such as sharp peaks of hypersensitivity and footprints, that result from protein/DNA interactions.

## Materials and Methods

### Sequence bias correction

Enzymes that are commonly used in molecular biology have nucleic acid preferences for their substrates and the sequence at or near the active site of the enzyme typically dictates enzymatic preference. Let the sequence context be defined by a k-mer proximal to the start of the detected sequence read. A sequenced read corresponds to a k-mer observation if it occurs at a specific offset with respect to the edge of the k-mer (Figure 1A). Assuming a systematic k-mer dependent bias, the true read count will be a scaled version of the observed read count, that is:

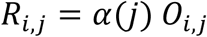

where, for position *i*, *R*_*i,j*_ is the true read count, *O*_*i,j*_ is the observed read count and *α*(*j)* is the scale factor, or bias, corresponding to k-mer *j*.

**Figure 1.**
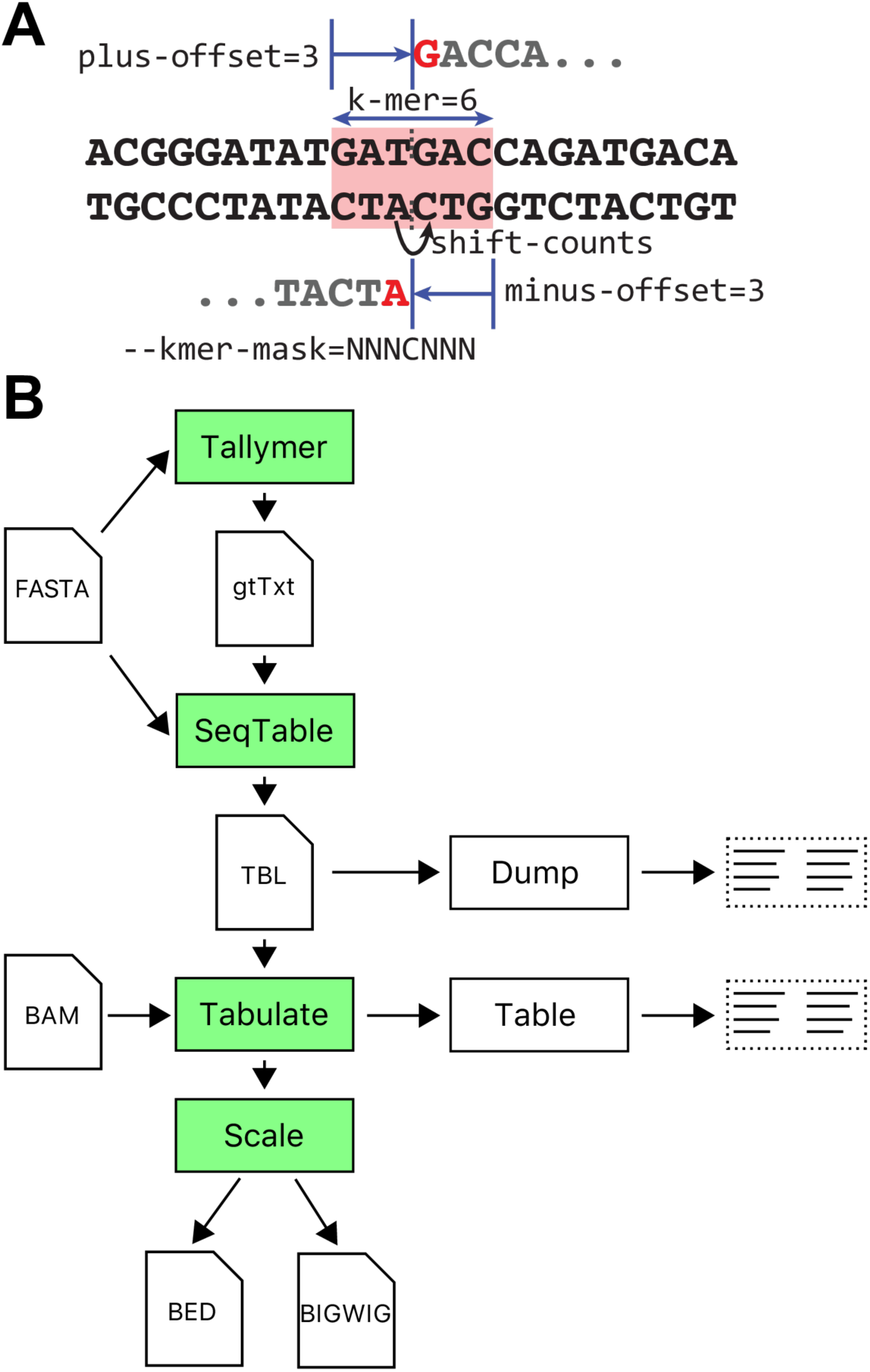
*SeqOutBias* overview and parameter definitions. A) An enzymatic cleavage event that results in a blunt end can be detected by sequencing the upstream or downstream DNA (red bases). The hexamer sequence centered (red block) on the nick sites (dotted vertical lines) confers specificity; this parameter is referred to as the k-mer. The plus-offset and minus-offset parameters specify the nick site relative to the first position and last position of the k-mer. As opposed to specifying the immediate upstream base for the minus strand, we shift the base position by +1 to match the first position of the plus aligned read. B) This panel illustrates the high-level overview of the inputs, intermediate files, and output of the *seqOutBias* program and the computation steps that the program performs. The *tallymer* step indexes the reference sequence (FASTA) and computes mappability for the given read length. The *seqTable* step parses the reference sequence together with the mappability information to compute the k-mer that corresponds to each possible read alignment position. The *tabulate* step tallies the k-mer counts across the selected regions (or the full genome), as well as the k-mers corresponding to observed aligned reads (if a BAM file is supplied). Lastly, *scale* computes the genome-wide aligned read pile-ups, scaling sequence reads by the expected/observed k-mer frequency.

In an unbiased setting, the observed frequency of k-mers should be proportional to their genome-wide counts or a regional subset of the genome. For example, condensed chromatin may restrict enzyme access and these can be excluded from the genomic k-mer counts. Let *I(m,j)* be the indicator function that genome position *m* is assigned to k-mer *j*, then we have:

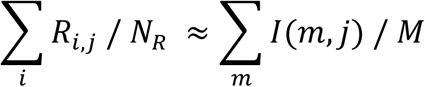

where *M* is the total number of observable genome positions and *N*_*R*_ is the total read count. Further taking the observed total read count *N*_*0*_ to be *N*_*0*_ *≈ N*_*R*_, then *α*(*j)* can be approximated by:

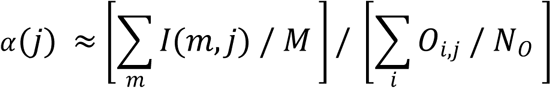

The *seqOutBias* software aims to correct sequence biases by scaling the aligned read counts by *α*(*j),* which is effectively the ratio of genome-wide expected read counts to the observed sequence counts for each k-mer. The *seqOutBias* software additionally takes into account mappability, which means that observable positions can differ per strand, thus we compute a separate value of *α*(*j)* on each strand.

### Computational workflow of seqOutBias

The *seqOutBias* software uses a genome FASTA file and an aligned and sorted BAM file as inputs. Each intermediate step within *seqOutBias* corresponds to a *seqOutBias* subcommand, which permits them to be run individually or separately on different machines. Due to the large size of genomic datasets, *seqOutBias* can read compressed FASTA files.

#### 1) Computing genomic mappability

In the implementation of *seqOutBias*, our algorithm first calculates the expected sequence detection frequency for each k-mer by determining the positions in the supplied reference genome that are uniquely mappable for a given sequence read length. It is important to compute mappability for a read length because the algorithm should only count k-mers that correspond to genomic positions that have the potential to be detected based on their unique genomic mappability when it calculates the expected k-mer detection frequency from the FASTA file. *SeqOutBias* invokes GenomeTools’ *tallymer* to compute mappability at each position in the genome (17, 18). First, the reference genome FASTA file is indexed using GenomeTools’ Tallymer program (17, 18). Next, Tallymer computes unique mappability for each position in the genome for a given input read length. This step corresponds to a *seqOutBias* subcommand, *tallymer*, which can be run individually or on different machines. The tallymer subcommand computes the mappability information, parses the reference sequence to compute k-mer indexes, and creates a mappability file for a given read length (Figure 1B). This process consists of three parts: 1) creating a suffix tree; 2) creating a genome index; and 3) creating the mappability file. These processes are the most computationally intensive steps and *seqOutBias* will recognize the existence of intermediate files in the directory to avoid unnecessary recomputation. For instance, if the *seqOutBias tallymer* step is executed for different read lengths, but using the same FASTA file, then the first suffix-tree portion is re-used across invocations.

#### 2) Mapping k-mer indexes to the aligned read positions

The next step, *seqOutBias seqtable*, creates an intermediate table that combines mappability information, read length, and plus/minus offsets (Figure 1A) to map k-mer indexes to the aligned read positions (Figure 1B). The *seqtable* subcommand parses the reference sequence (FASTA) together with the mappability information to compute the k-mer that corresponds to each possible read alignment position. The resulting binary file stores this information in a compressed form that can be easily used for subsequent computation steps, as well as storing the corresponding parameters (read length, k-mer size, and cut-site offsets). This intermediate file reduces the amount of computation needed when processing aligned read files and provides an intermediate TBL file that decouples the reference sequence processing from the remaining steps.

#### 3) Tallying the k-mer counts in the reference sequence and the aligned reads

The resultant TBL file is an input for the *seqOutBias tabulate* subcommand, which tallies the k-mer counts across the selected regions (or full genome), as well as the k-mers corresponding to observed aligned reads from the BAM file. In contrast to other methods (10–12, 15), these numbers are used to scale the reads without the need for Naked DNA to calibrate. This subcommand produces a k-mer count table based on the TBL sequence information and the optional sorted BAM file. Counts correspond to the entire genome by default, but counts can be constrained to specific regions by supplying a BED file with the *regions* option. When no BAM file is supplied, the output will have four columns: k-mer index, k-mer string, plus strand count, and minus strand count. If a BAM file is supplied, the output will have two additional columns with the plus and minus strand counts of observed aligned reads.

#### 4) Scaling individual sequence reads

The final subcommand, *seqOutBias scale*, computes the genome-wide aligned read pile-ups and scales them by the expected/observed k-mer detection frequency. This produces the corrected aligned read pile-ups, both as BED and bigWig files. This command provides flexibility in the output, including the *--shift-counts and* --*tail-edge options.* The *--shift-counts* option shifts minus strand pile-up positions to align with the plus strand pile-up, making reads from both sides of a cleavage site pile up at the same position regardless of whether the upstream or downstream sequence was detected by sequencing (Figure 1A). This option is used when enzymatic cleavage of individual sites can result in a single base shift depending on whether the nicking event was detected by sequencing the upstream or downstream DNA (red nucleotides in Figure 1A). The *tail-edge* option outputs the 3´ end of the reads; this option is used primarily for analysis of PRO-seq data (4, 5). Therefore, *seqOutBias* reads compressed files (FASTA, mappability information, and sorted BAM files), reuses intermediate results, and allows for flexibility in specifying sequence features for data correction.

### k-mer mask optimization

In *seqOutBias*, the k-mer sequence that is recognized by the enzyme to confer specificity is characterized by three parameters: k-mer size and a pair of offsets for the plus and minus strands (Figure 1A). These parameters enable flexibility and *seqOutBias* works with enzymes that have a variety of recognition site lengths. The *k-mer* mask parameter of *seqOutBias* restricts which positions contribute to the bias correction. A gapped k-mer enables the user to capture bias due to distal contributions and ignore uninformative positions that are more proximal. Positions within the mask that are ignored are represented by an *X* and informative positions by an *N*. This parameter provides an alternative way to specify the position intervening between the first base sequenced and the base directly upstream by inserting a *C* in the mask string. For example, a possible 8-mer that spans 16 bp could be represented as *NNXXNNXXCXXNNXXNN*; likewise, *NNNCNNN*, would represent a recognition site with *kmer-size* = 6, *plus-offset* = 3 and *minus-offset* = 3 (Figure 1A). We developed two approaches to guide the choice of the k-mer mask.

The first of these approaches, implemented in the Rust program *kmer_mask_em* (<u>https://github.com/guertinlab/kmer_mask_em</u>), is aimed at assays like DNase-seq, where an enzyme cuts with a preferred orientation and this cleavage event can be detected by sequencing the upstream or downstream sequence. The second approach, implemented in R (<u>https://github.com/guertinlab/seqOutBias/tree/master/docs/R</u>), aims to flatten the composite profiles of the input molecular genomics data at transcription factor position specific weight matrices (PSWM) and is offered as an alternative that imposes less constraints on the k-mer mask.

#### 1. k-mer mask optimization via enzyme cut-site model

In the k-mer mask optimization via enzyme cut-site model method, we model the enzyme cut bias as a PSWM of length K, over the sequence surrounding the cut site. Each k-mer corresponds to a possible cut-site with unknown orientation. We apply expectation maximization (EM) to infer the PSWM from the table of k-mer counts.

Let the PSWM be defined as *θ*= {*p*_*j,b*_ *: a≤ j ≤ K; b ∈ {A, C, T, G}}*, where *p*_*i,b*_ be the probability of observing the nucleotide *b* at cut-site position *j* and 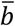 to be the complement of nucleotide 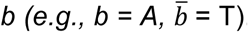. Given a table of k-mer counts, let *x*_*i*_ be the sequence of k-mer *i* occurring *n*_*i*_ times, *z*_*i ∈*_ *{ fwd, rev }* be unknown orientation of *x*_*i*_ and *P(Z*_*i*_ *= fwd) = γ.* The full likelihood of the model is depicted in Figure S1, thus the full log-likelihood is given by:

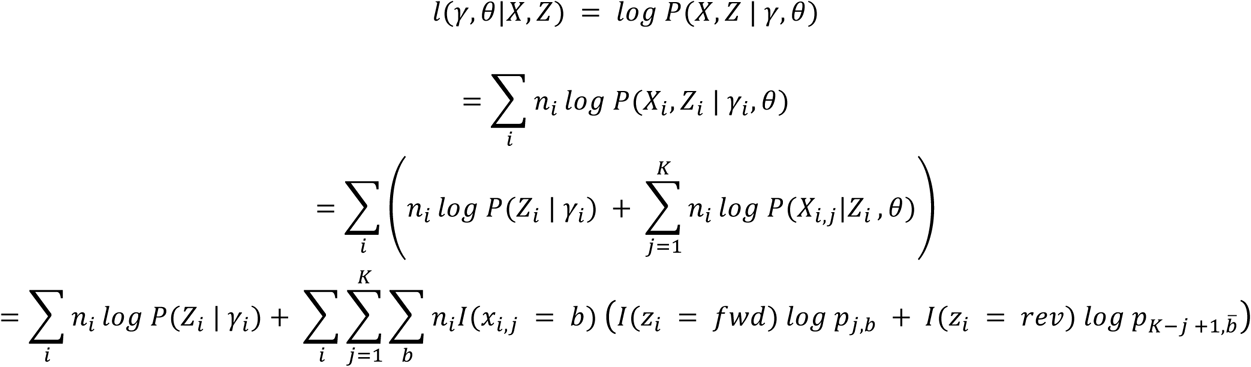

Which results in the simple definitions for the EM steps:

E-step: use Bayes rule to get the posterior of Z, i.e., compute *P (Z*_*i*_ *= fwd | X*_*i γ*_ *, θ);*

M-step: update the model parameters:

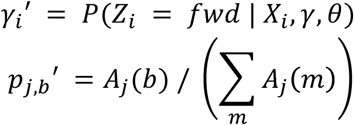

with:

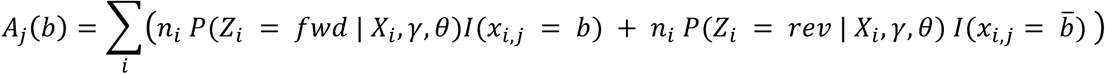

In practice, we add a pseudo-count to the values of *A*_*j*_*(b)* of 0.1. Furthermore, since EM does not guarantee a global optimum, the *kmer_mask_em* program computes the optimization from multiple random starting points, taking the best global result as the optimum.

The inferred PSWM is used to infer a sequence of k-mer masks of increasing complexity by making use of the Information Content score at each position. Positional Information Content refers to the relative difference in observed sequence at that position relative to what is expected, so high Information Content means that a position is likely to influence enzyme preference. Bases that have a value lower than a given Information Content threshold are excluded from the mask. Finally, the resulting “forward” mask is combined with it’s own “reverse” mask. Therefore, a position in k-mer mask is unmasked if it is unmasked in either the forward or the reversed mask. This is necessary since during the execution of the *seqOutBias* program we do not know the orientation of each specific site.

This approach is ideal for assays where a single enzyme cuts with a preferred orientation, producing reads in both directions. The resulting matrix provides candidate positions that are influencing the enzyme specificity. This approach: 1) assumes that the mask is symmetric; 2) requires a full counts table (all positions unmasked) as input; and 3) requires multiple runs of the same computation with random starting sites, which are automatically done in parallel, to ensure a reasonably good global optimum for the PSWM.

#### 2) k-mer mask optimization using profiles at PSWMs and hill climbing optimization

We implemented a hill climbing method to optimize the k-mer mask. This method takes a starting k-mer mask and a set of read count tables (one for each transcription factor) as inputs to guide a greedy search over the space of possible k-mer masks. The metric we use to evaluate k-mer masks aims to measure the effect these masks have on the TF composite profile at the binding sites.

For k-mer mask *m* and a given TF *t*, let *c*_*i,j*_ *(t,m)* be the scaled read count at position *j* of the binding site *i*, then the TF profile is the vector *G*_*t*_*(m) = [p*_*j*_*(t,m)],* where:

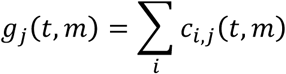

Our metric is then defined as:

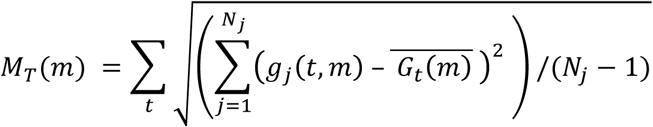

Let *H(m)* be the set of masks that differ from *m* by only one *X* being changed to *N* (i.e. by unmasking an additional position). The search procedure, given a starting mask *m*_*0*_ (for example the mask of all *X*, indicating that all positions in the mask are excluded from the k-mer), is simply the iteration:

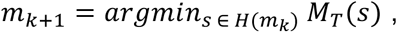

stopping when there are no more masked *X* positions in the current mask.

This empirical approach requires many k-mer mask evaluations, which correspond to complete runs of *seqOutBias*. Our implementation allows multiple instances of *seqOutBias* can be run in parallel (see mc.cores parameter in <u>https://raw.githubusercontent.com/guertinlab/seqOutBias/master/docs/R/seqOutBias_hcsearch.R).</u>

### SeqOutBias Application Programming Interface (API)

We structured the code into two parts: a main library (*seqoutbiaslib*) and the command line program (*seqOutBias*) implemented using the main library. This split allows the code to be reused to implement different interfaces with similar functionality or as a component in a larger program. The library is exposed as both a Rust library and a C library.

The Rust interface includes everything used to build the main program. The code is split into a series of modules which correspond to the subcommands of *seqOutBias*:

- *tallyrun* – code to execute GenomeTools (17) to produce the mappability file
- *tallyread* – code to read and access the mappability information
- *seqtable* – code to read and write seqtable files to disk
- *fasta* – code to read in the FASTA file, combine it with the mappability information, and produce the seqtable file (via calls to the seqtable module)
- *filter* – code to filter BAM records based on things like length, quality, etc.
- *counts* – code to tabulate kmer counts
- *scale* – code to compute read pile-ups and scale them appropriately
- *bigwig* – code to write the chromInfo and wiggle files and convert them to a bigWig file using the wigToBigWig program

The C API exposes the ability to generate a seqtable file and query pile-ups in memory without the need to write them to disk. The C API exposes two query functions, one to query specific genomic coordinates and one that returns a full chromosome as an array. The C library and the corresponding header files are built as part of the main compilation process and can be linked as typical for C libraries. Functions can be grouped into four sets:

1. Functions to manage the *seqtable* generation parameters.
2. A function to create a default set of pile-up generation parameters.
3. A function to generate the seqtable file.
4. Functions to generate and query the pile-ups.

### Deproteinized DNA ATAC-seq

The naked DNA ATAC-seq library was prepared as previously described (7) with several modifications: 1) we used purified genomic DNA, as opposed to crude nuclei isolations; 2) we omitted IGEPAL CA-630 from all buffers; and 3) we performed PCR cleanup using AMPure XP beads to select DNA <600 bp. The naked DNA ATAC-seq data were deposited in the Gene Expression Omnibus (GEO) database, with accession number GSE92674.

### Installation and analyses

The user guide and install instructions are available through GitHub: <u>https://guertinlab.github.io/seqOutBias/seqOutBias_user_guide.pdf.</u>

The analyses presented herein are reproduced in full with rationale in the accompanying *seqOutBias* PDF vignette on GitHub: <u>https://guertinlab.github.io/seqOutBias/seqOutBias_vignette.pdf</u>. We also provide a website version of the vignette: <u>https://guertinlab.github.io/seqOutBias_Vignette/.</u>

## Results

### Correction of individual DNase-seq reads

DNase-seq measures the accessibility of the phosphodiester backbone of DNA at single-nucleotide resolution (1, 2, 9, 19). Composite DNase-seq profiles that are centered on sequence motifs of TF binding sites accentuate molecular features that inform on TF binding properties. For example, DNase footprints are defined as depletions of sensitivity within large regions of hypersensitivity; footprints align with TF recognition sites and result from TF interactions with DNA (20, 21). High throughput DNase-seq experiments described a cleavage pattern at the footprint that was interpreted as a measure of TF/DNA interactions (9); however, subsequent work attributed these artifactual signatures to differential substrate specificity of DNase conferred by the presence of the TF motif (10–12). As a result, some footprint detection programs now incorporate sequence biases into their algorithms (12, 15, 22). *SeqOutBias* provides the option to correct enzymatic sequence bias prior to footprint detection and the output files can be used with existing footprinting algorithms that do not incorporate a correction step.

Previous studies used a hexamer (11) and a tetramer (10) centered on the DNase cut site to account for the intrinsic sequence bias of DNase. We systematically explored how the individual bases within a 10-mer contribute to the preference of DNase (see Methods). We found that there was little Information Content beyond position 3 from the DNase cut site (Figure S2A&B). This expectation maximization method identifies candidate positions that contribute to enzyme specificity based on sequence content, so we sought to directly test how each position in the mask contributes to the smoothing of the composite profiles. For each TF PSWM, we computed the standard deviation for the profile obtained by summing the scaled reads across all sites at each position in the PSWM (see Methods). We summed these standard deviations across a set of PSWMs as a metric to determine the contribution of each position to DNase preference. We found only a modest improvement beyond tetramer correction (Figure S2C).

We scaled individual reads based on the preference of DNase using *seqOutBias* and a 6-mer correction factor (Figure 2). Figure 2A illustrates that DNase prefers to nick the sequence *CCTTGC* and the read associated with this window was reduced to an intensity of 0.15. DNase disfavors nicking of *GGGGAA*, thus the read associated with this hexamer was scaled to an intensity of 5.6.

**Figure 2.**
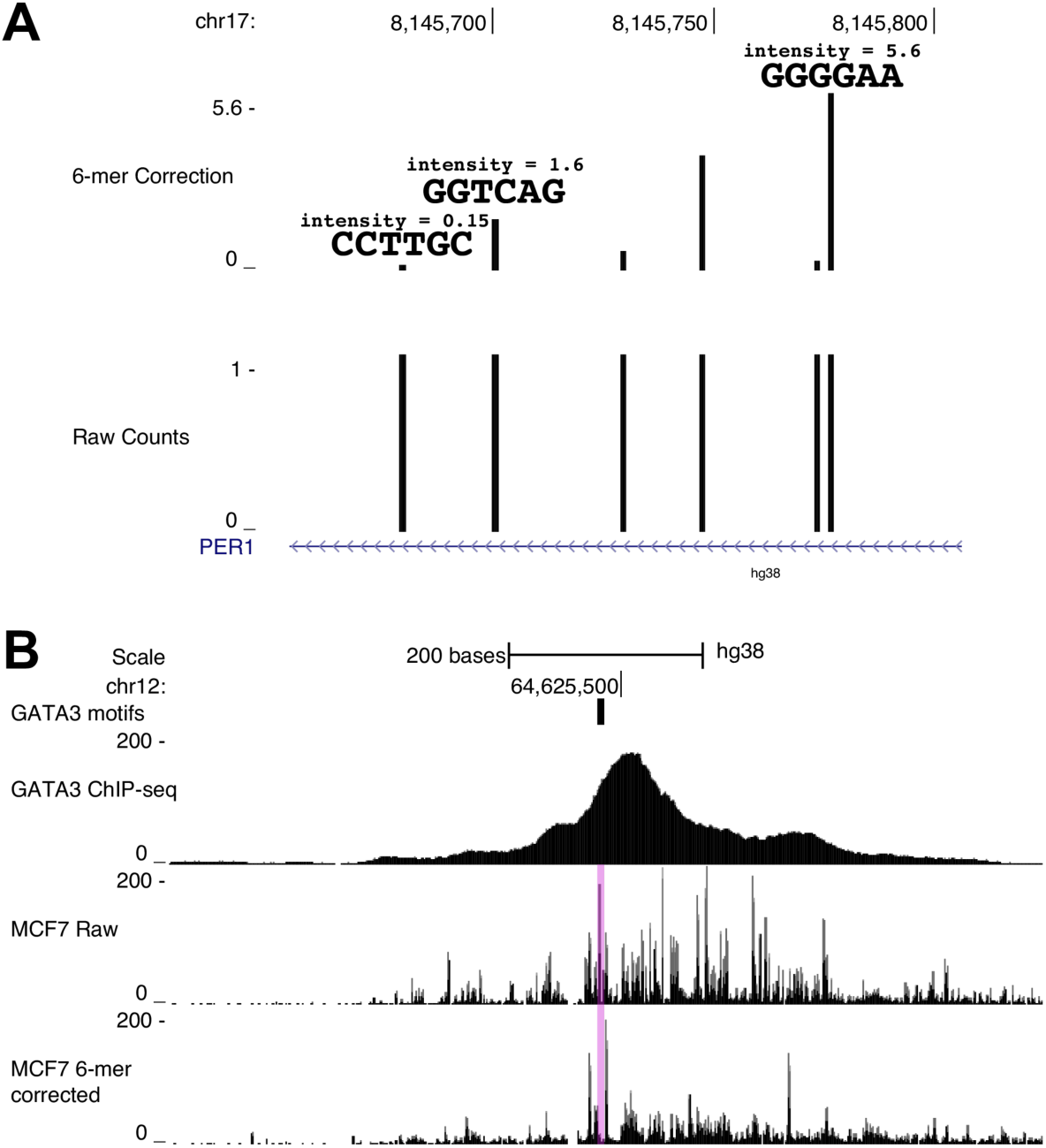
*SeqOutBias* scales individual sequence reads and corrected DNase-seq data reveals footprints. A) The bottom track shows six nick positions from naked DNA DNase-seq data; each position was found once in the data. The top track reports corrected read intensities, which scale inversely with DNase sequence preference. B) The GATA3 binding site (transparent pink) contains sharp peaks within the binding site in uncorrected DNase-seq profiles; a footprint is present only in the corrected data.

DNase sequence preferences are most apparent in composite profiles of DNase cut frequency surrounding TF motifs. We tested the efficacy of 6-mer correction on DNase-digested naked DNA (23); corrected profiles of naked DNA digestion should not exhibit footprints or molecular signatures that result from protein/DNA interactions. We observe that sharp peaks and troughs are smoothed in the corrected composite profiles for ELF1, GATA3, and MAX motifs (Figure 3).

**Figure 3.**
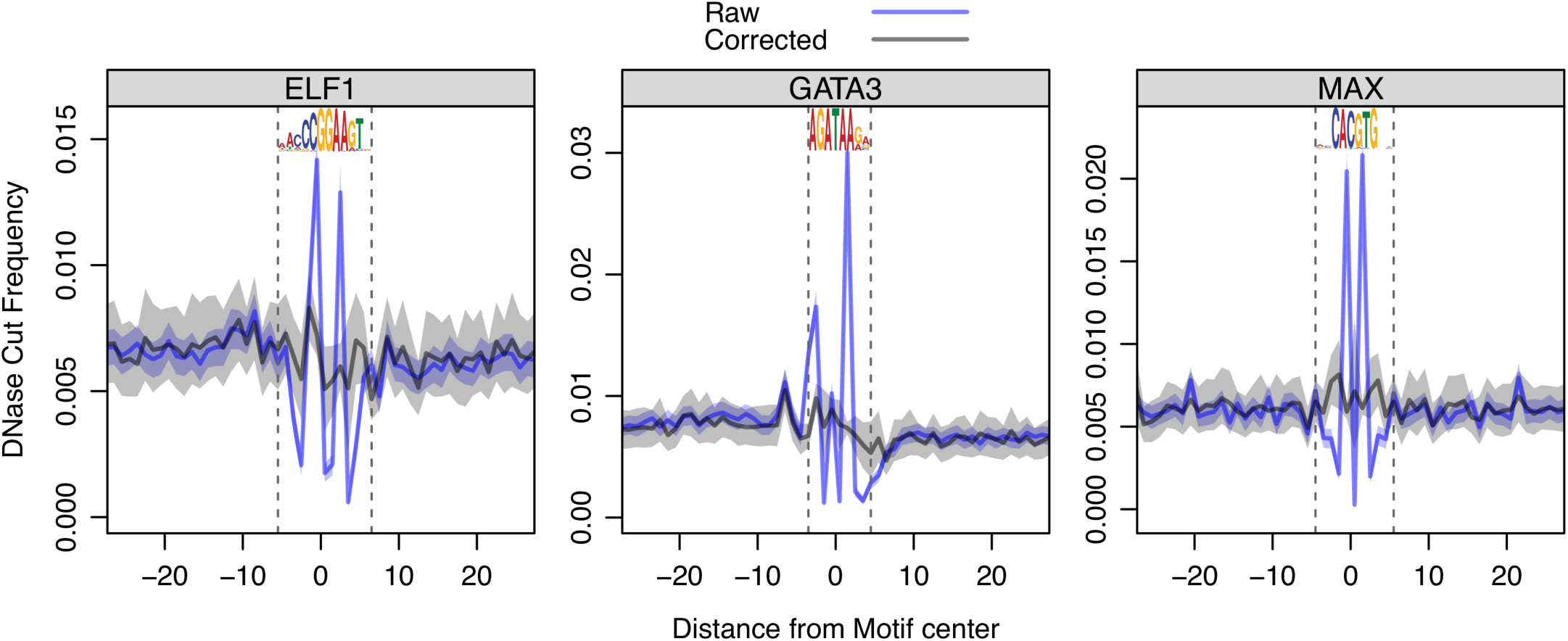
DNase nick bias is corrected in a naked DNA DNase experiment. Each composite profile illustrates the average cut frequency at each position between nucleotides. The blue trace is the raw data and the black trace is the corrected data; the opaque boundaries represent the 75% confidence interval. A seqLogo representation for each TF’s binding site is shown at the top of each plot and vertical dashed lines show the boundaries of sequence information content.

True signatures that result from TF/DNA interactions are not smoothed by *seqOutBias*. For instance, ChIP-seq validated CCCTC-binding factor (CTCF) binding sites exhibit strong footprints and composite profiles at CTCF motifs highlight a sharp signature upstream of CTCF binding (24). This CTCF signature is unaffected after correcting for DNase intrinsic sequence preference (12). We plotted DNase-seq profiles at GATA3 and MAX binding sites to determine whether true molecular signatures are apparent after intrinsic bias correction (Figure 4). We observe a clear composite footprint at MAX binding sites in chromatin, as expected, this footprint is not present in the naked DNA digestion (Figure 4A). The MAX footprint is obscured by sharp peaks of hypersensitivity (sequence artifact signatures) in the raw uncorrected traces (Figure 4A). Individual footprints also exhibit these sharp peaks at the site of TF binding in raw data. At a ChIP-seq validated MAX binding site we observe a footprint only after correcting for DNase sequence bias (Figure S3). We observe a sharp DNase signature upstream of GATA3 binding sites, which is present only in the chromatin digested samples (Figure 4B). We conclude that this molecular signature is a result of GATA3/DNA interactions, because this peak is neither smoothed following *seqOutBias* correction nor present in the naked DNA DNase digested sample. Note that GATA3 does not have an appreciable composite footprint, but TF inference algorithms may use TF-specific signatures, as we observe for GATA3, to inform on TF occupancy and binding intensity. The sharp peak with raw DNase-seq data within the GATA3 motif obscures footprints at individual GATA3 binding sites (Figure 2B and Figure S4). Bias correction enhances both the footprint and the signature upstream of the GATA motif (Figure S4). Therefore, correction of intrinsic DNase sequence bias highlights true molecular features: footprints and sharp hypersensitivity peaks. We propose that these features can be systematically characterized for all TFs and used as informative priors when inferring TF binding profiles genome-wide from enzymatic hypersensitivity data.

**Figure 4.**
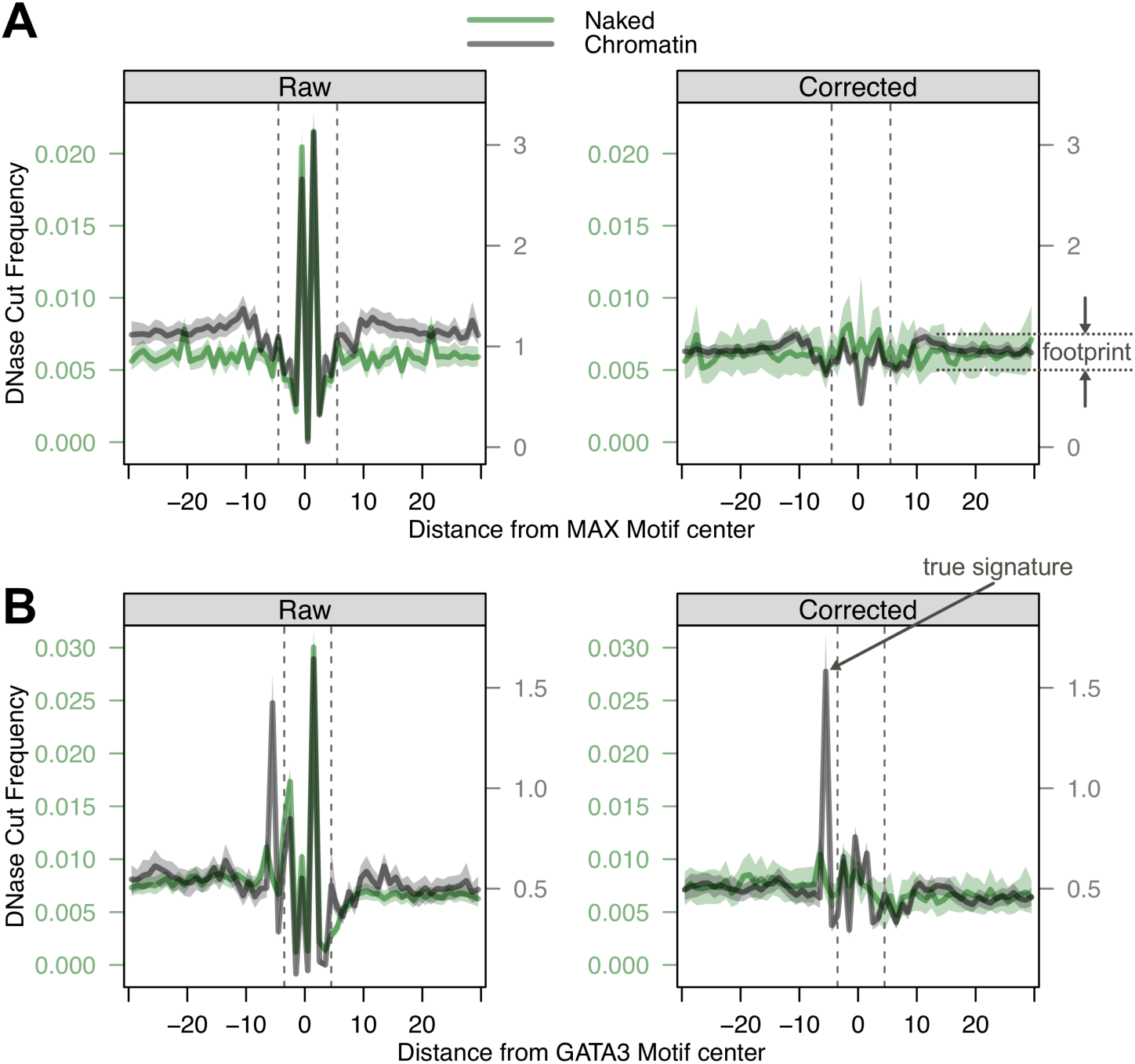
True molecular signatures resulting from TF/DNA interactions are visible in corrected composite profiles. A) A true footprint is highlighted in corrected composite profile (right panel) of DNase cleavage at ChIP-seq confirmed MAX binding sites (29) compared to raw frequency counts (left panel). The black trace is DNase-digested chromatin and the green trace is DNase-digested naked DNA. As expected, the composite footprint is not detected in the naked DNA composite. B) A true molecular signature is highlighted in the corrected composite profile (right panel) of GATA3 binding sites (29). The signature is exclusively detected in the chromatin digested experiment and may result from GATA3/DNA interaction.

### Correction of TACh-seq, MNase-seq, ATAC-seq, and PRO-seq data

We characterized and corrected the biases of Benzonase and Cyanase using Tissue Accessible Chromatin (TACh-seq) data (6). TACh-seq is a variant of traditional enzymatic hypersensitivity assays whereby frozen tissue samples are treated with either Benzonase or Cyanase endonuclease. Benzonase is an endonuclease cloned from *Serratia marcescens* that functions as a dimer and Cyanase is a non*-Serratia* monomeric enzyme. These enzymes are more highly active under high salt and high detergent conditions, so these enzymes are more suited for digestion of solid tissue sample, which requires harsh dissociation treatments. We corrected TACh-seq data generated from frozen mouse liver tissue (6). Composite profiles from CEBP-beta, FOXA2, and CTCF binding sites (25–27) in mouse liver indicate that an eight base pair mask centered on the nick site is sufficient to correct both Cyanase and Benzonase biases (Figure S5 and Figure S6). Next, we applied *seqOutBias* correction to MNase-seq data generated from MCF-7 cells (28). An eight base pair mask abrogates the intrinsic sequence bias of MNase-seq data (Figure S7).

ATAC-seq is unique among enzymatic accessibility assays because each transposition event inserts two sequencing adapters into the chromatin. Each Tn5 molecule can be pre-loaded with any combination of the paired-end 1 and paired-end 2 adapter. Reads that align to the plus and minus strand are processed separately because the Tn5 recognition site is distinct for plus and minus reads. We applied *seqOutBias* correction to published ATAC-seq data from GM12878 cells (7). We generated and analyzed naked DNA libraries using the ATAC-seq work flow to measure Tn5 specificity in the absence of chromatin (GEO accession: GSE92674). We optimized the k-mer mask for ATAC-seq data by starting with a k-mer mask of 12 *X* bases flanking each side of the Tn5 insertion site, then we systematically changed each *X* position into a masked *N* using a hill climbing mask optimization method (see Methods). We chose the position that results in the lowest summed standard deviations across a set of PSWMs and iterated until we found the top 11 positions that contribute to Tn5 sequence bias (Figure S8). We used the *N* positions of the 8-mer *NXNXXXCXXNNXNNNXXN* for the ATAC mask because these are the most influential for Tn5 recognition of plus strand reads (Figure S8). The reciprocal mask, *NXXNNNXNNXXCXXXNXN*, is the optimal 8-mer mask for minus strand reads. The sharp ATAC-seq spikes at the sites of TF binding for SP1, REST, and EBF1 (29) are reduced in the corrected data (Figure S9 and Figure S10). The complex nature of Tn5 recognition and dual loading of adapters, taken together with the incomplete smoothing of ATAC composite profiles, suggests that a simple spaced k-mer correction may not be sufficient to fully correct Tn5 bias.

PRO-seq couples terminating nuclear run-on assays with high throughput sequencing to quantify engaged RNA polymerase molecules genome-wide at nucleotide resolution (4). Sequence composition of transcripts may affect run on efficiency, therefore, the sequence immediately downstream of RNA polymerase may influence detection of RNA molecules. The sequence upstream of RNA polymerase could affect ligation efficiency because T4 RNA ligase treatment may exhibit sequence preference. We used *seqOutBias* to scale published PRO-seq data from K562 cells (30). We specifically used annotated transcripts to calculate expected k-mer frequency, as opposed to genomic k-mer frequency, because the vast majority of transcription occurs within gene annotations (31). We found that a k-mer mask that spans the last three bases of the ligated RNA molecule and the three bases downstream from RNA polymerase is sufficient to correct the PRO-seq data (Figure 5 and Figure 6). RNA polymerase density decreases at the polypyrimidine tract upstream of the 3´ splice site, which suggests an increased RNA Polymerase elongation rate at this tract (Figure 6). These data indicate that in addition to U2AF (32, 33) recognizing the pyrimidine residues in the pre-mRNA polypyrimidine tract, this tract increases RNA polymerase elongation rate.

**Figure 5.**
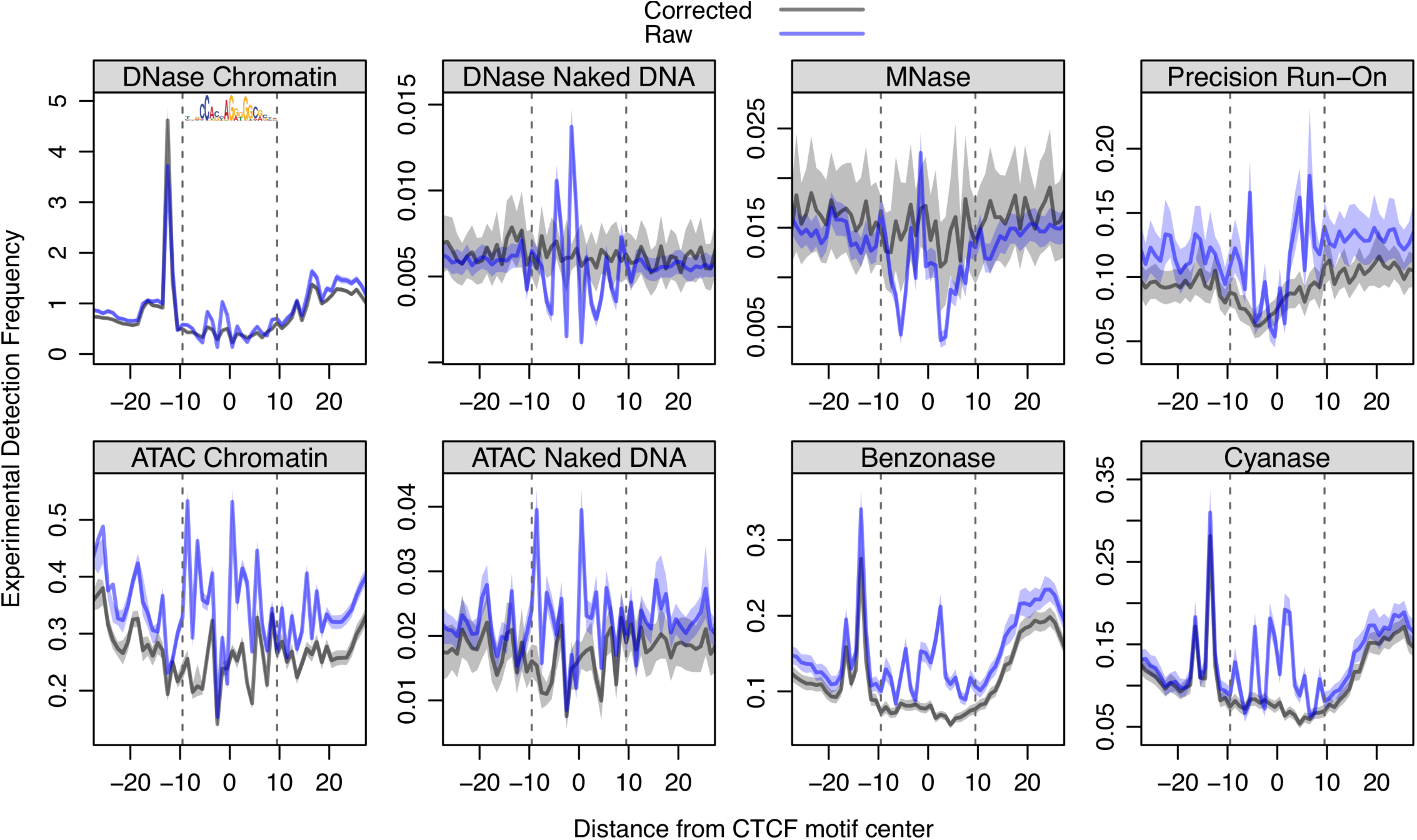
*SeqOutBias* corrects sequence bias at CTCF binding sites associated with DNase (9, 23), Tn5 Transposase (ATAC) (7), Benzonase (TACh) (6), Cyanase (TACh) (6), MNase (28), and T4 RNA ligase (PRO) (4). Upon correcting for enzymatic sequence bias, the artifactual spikes at the CTCF binding site are abrogated in each molecular genomics dataset we tested. However, in cases of CTCF binding to chromatin, we observe protection that results in a footprint; note that MNase is not expected to result in a composite footprint. We observe the previously characterized sharp peak upstream of the CTCF motif and this molecular signature is likely caused by CTCF-mediated enhancement of cleavage activity.

We also observe a sharp peak at position −3 from the 5´ end of exon. This sharp peak is absent using *seqOutBias*-corrected reads (Figure 6), therefore we hypothesized that this peak results from either inefficient adenine incorporation during the nuclear run-on or a preference for cytosine or uracil during either the run-on or ligation reaction. We classified 3´ splice acceptor sequences into AAG, CAG, and TAG to generate composite RNA polymerase profiles--note that very few splice acceptor sequences are GAG, so they were excluded. The sharp peak at position −3 is exclusively found in the CAG splice acceptor profile (Figure S11), which indicates that cytosine is preferentially incorporated during the run-on or preferentially ligated. We examined the composite profiles at corrected CAG and TAG splice acceptor sites and we observed that RNA polymerase density is higher following CAG splice acceptor sites (Figure S12). This result indicates that the consensus splice acceptor site, CAG, decreases RNA polymerase elongation rate in the 5´ region of the exon.

**Figure 6.**
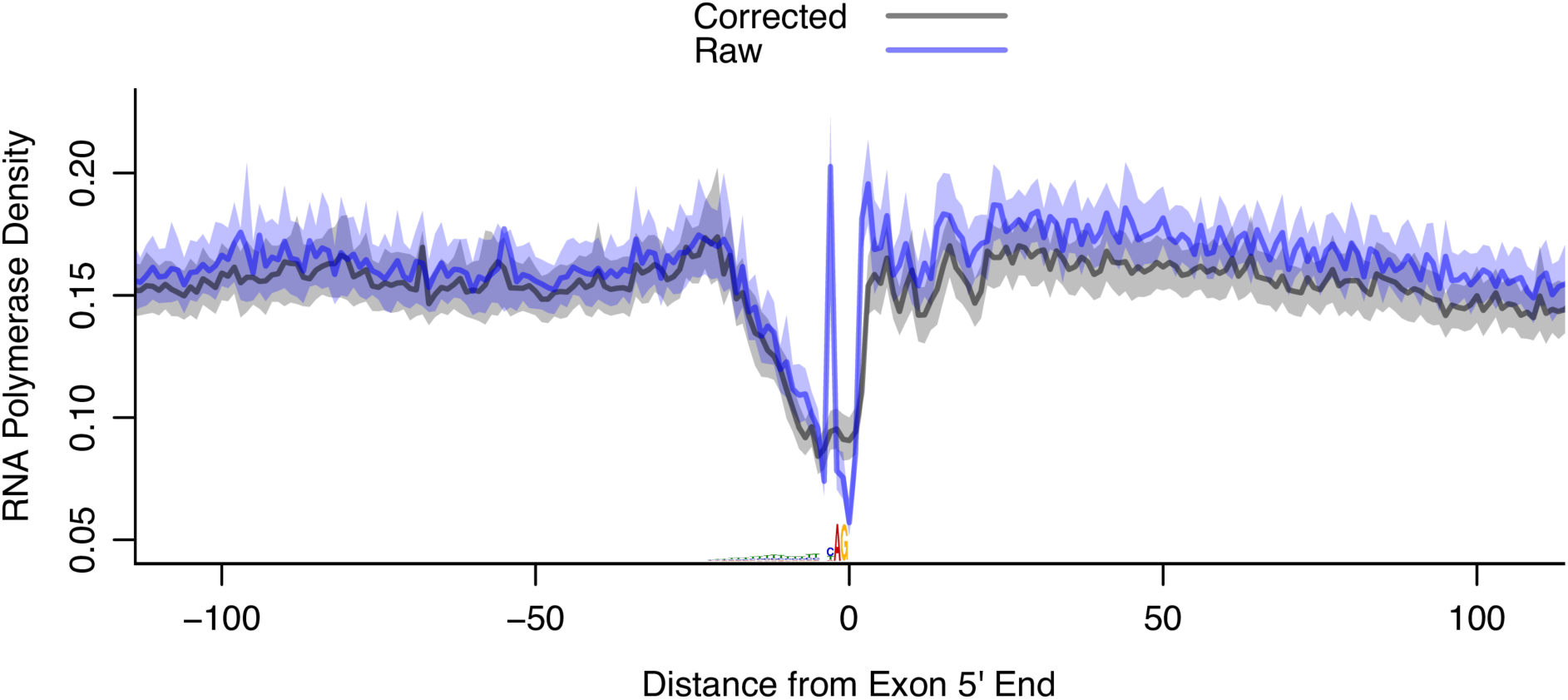
*SeqOutBias* corrects sequence bias associated with the 3´ splice site recognition motif. Upon correcting for enzymatic sequence bias, the artifactual signature at the 3´ splice site is abrogated. The first base of the exon spans position 0-1 on the x-axis and the sequence bias peak is at position −3.

Genome-wide binding data for CTCF is available for K562, GM12878, mouse liver, and MCF-7 cells (27, 29). Upon correcting for enzymatic sequence bias, the sharp signature artifacts at CTCF motifs are abrogated in each molecular genomics dataset we tested (Figure 5). The naked DNA profiles for ATAC-seq and DNase-seq are not restricted to CTCF-bound sites; all genomic CTCF motifs are included in these composites (Figure 5). In the chromatin TACh, DNase, and ATAC experiments, we observed protection resulting in a footprint and a sharp peak upstream of the CTCF motif. Taken together, we show that *seqOutBias* effectively corrects enzymatic sequence bias resulting from a diverse set of molecular genomics experiments.

### Enzymatic DNA end repair and ligation bias

We found that the bases upstream and downstream of a DNase nick site are not equally likely to be detected by sequencing (red nucleotides in Figure 1A). In Figure 1A, for *GATGTC* we would expect the ratio of reads that begin with *GACCAGATGACA* (plus strand) and *ATCATATCCCGT* (minus strand) to be approximately equal to one if this site was nicked repeatedly and there was no enzymatic end repair and ligation bias. We performed this analysis for all instances of each *NNNGAC*-mer in the genome (Figure 7A) and all 4096 pairwise combinations of 3-mers (Figure 7B). Palindromic 6-mers are balanced (Figure S13), but for most 3-mer combinations we identified a preference for which 3-mer is detected by sequencing, we term this “detection bias.” We took the top 5% most skewed 6-mers and generated a composite motif. This motif indicates that an Adenine in position 4 of the 6-mer is preferentially sequenced compared to the oppositely oriented nucleotide in position 3 (Figure S14).

**Figure 7.**
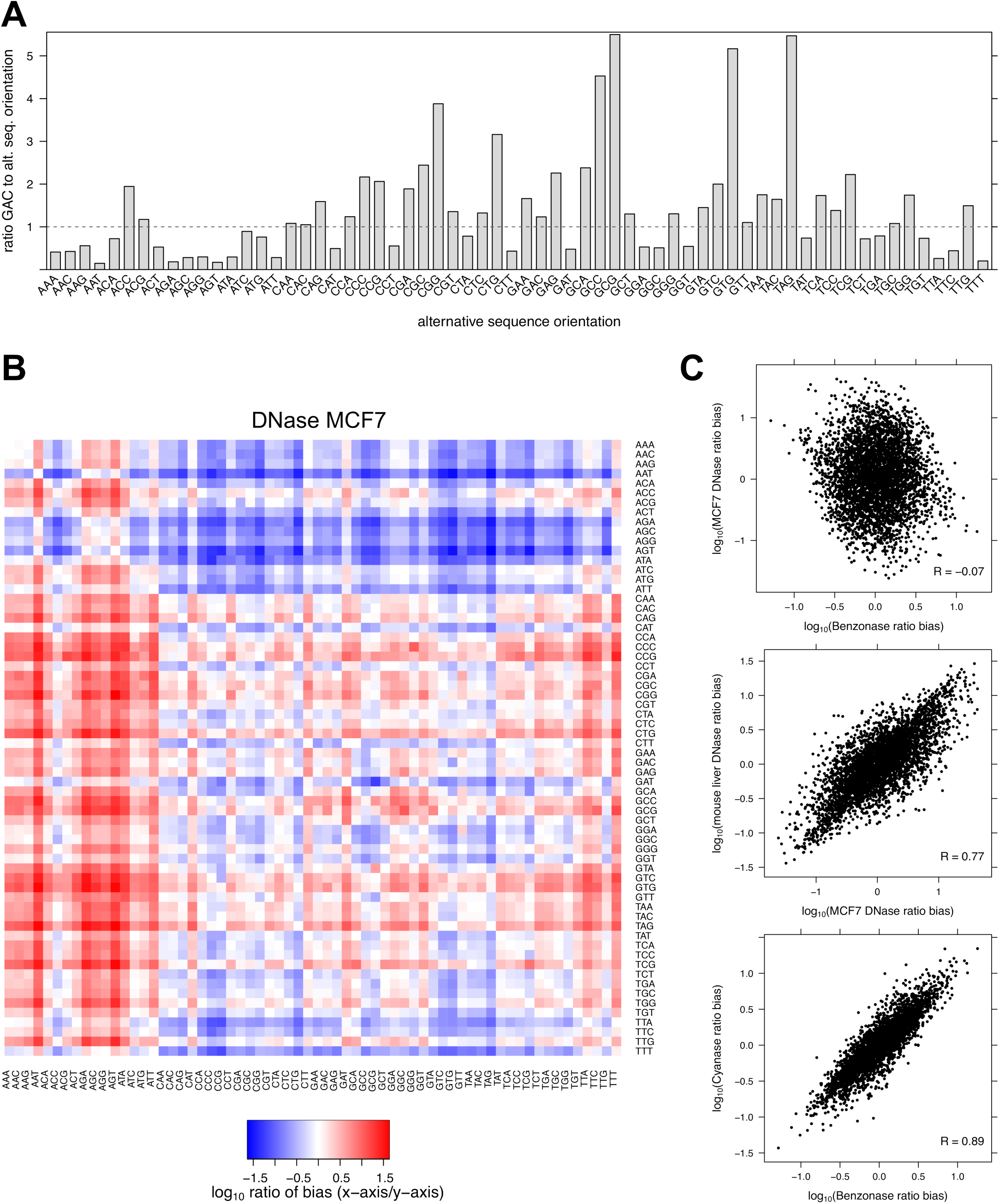
Detection biases are highly correlated between enzymes with similar cut specificities, suggesting that ssDNA overhangs drive enzymatic specificity. A) For all sequence-detected DNase-nicked 6-mers that end in *GAC* we compare the ratio of sequence reads that start with *GAC* to the oppositely oriented 3-mer. This bias results from enzymatic end repair and ligation sequence preference during the library preparation. B) The relative bias of all 3-mers sequenced (the ratio of x-axis 3-mer to y-axis 3-mer). Note that these k-mers are not clustered. C) This figure plots the values from panel B. The post-nick sequence preferences are highly correlated between DNase-seq experiments and between Benzonase and Cyanase experiments, but not between DNase and Benzonase.

Preparing digested DNA for Illumina high throughput sequencing requires several enzymatic treatments. T4 DNA Polymerase treatment removes 3´ overhangs and fills in 3´ recessed (5´ overhang) ends. T4 Polynucleotide kinase phosphorylates the 5´ end and Klenow Fragment (3´ to 5´ exo-) adds a single 3´ adenine overhang. We hypothesized that the overhanging sequences dictate the detection bias, because the detection bias is distinct for Benzonase and DNase (Figure 7C top panel). Although four nick events are necessary to sequence a DNA molecule, enzymatic hypersensitivity assays only detect one nick on each end of the molecule and it is impossible to determine the precise location of the other nicks. By assuming that two enzymes with similar nick specificity (Figure S15) will have comparable distribution of sequence overhangs, we can test the hypothesis that the overhang sequences contribute to post-nicking enzymatic treatment biases. We compared this post-nicking bias using DNase-seq data from two different labs and two different organisms (Figure S15). We also compared the detection bias of Cyanase and Benzonase, which have similar sequence preferences (Figure S15). Indeed, digestions with enzymes that have similar nick preferences, which results in comparable distributions of overhanging sequences, have highly correlated detection biases (Figure 7C bottom two panels and Figure S16). Importantly, *seqOutBias* calculates the ratio of genomic k-mers and experimentally observed k-mers to scale individual reads and this calculation inherently corrects for the convolution of biases resulting from multiple enzymatic steps, including these end repair and ligation biases.

## Discussion

We previously described the challenge of interpreting single-nucleotide resolution DNase-seq data (10, 11). Subsequently, groups have developed algorithms that consider this bias for DNase-seq footprinting detection (12, 22). However, this is the first report of stand-alone software that specializes in correcting sequence bias for a diverse set of molecular genomics datasets. *SeqOutBias* is a command line tool and designed for a UNIX environment, making the software compatible for seamless integration into existing high throughput sequencing analysis pipelines. *SeqOutBias* is conceptually and mathematically simple, effectively counting k-mer occurrences and scaling data accordingly. This calculation sufficiently corrects biases associated with many different assays. However, we anticipate that subsequent software may incorporate more complex calculations and models into data correction. For instance, RNA hairpins may affect the efficiency of ligating adapter to RNA using T4 RNA ligase. Due to the complexity of secondary and post-secondary RNA structure predictions (34), we suspect that more sophisticated models are necessary for correction of datasets such as PRO-seq.

Enzymatic hypersensitivity assays have the potential to identify regulatory elements genome-wide and infer TF binding intensity at each regulatory element. Four features of enzymatic hypersensitivity assays can aid in TF binding inference: 1) the presence of a TF’s recognition motif; 2) the raw enzyme cleavage frequency in the region surrounding the motif; 3) a depletion in sensitivity at the motif (footprint); and 4) the presence of TF-mediated molecular signatures (sharp peaks and valleys) that surround the motif. Correction of enzymatic sequence bias provides a more accurate measurement of all these features except sequence composition. Correction of intrinsic experimental biases will prove important as the field continues to refine experiments and algorithms to more accurately infer TF binding intensity genome-wide from enzymatic hypersensitivity data.

In conclusion, we and others have previously shown that enzymatic sequence preferences can be misinterpreted as biologically important phenomena (10–12). Sequence bias correction is an important step in analyzing high resolution molecular genomics data and we introduce *seqOutBias* as flexible and novel software that efficiently characterizes biases and appropriately scales individual sequence reads.

---

**Figure S1.**
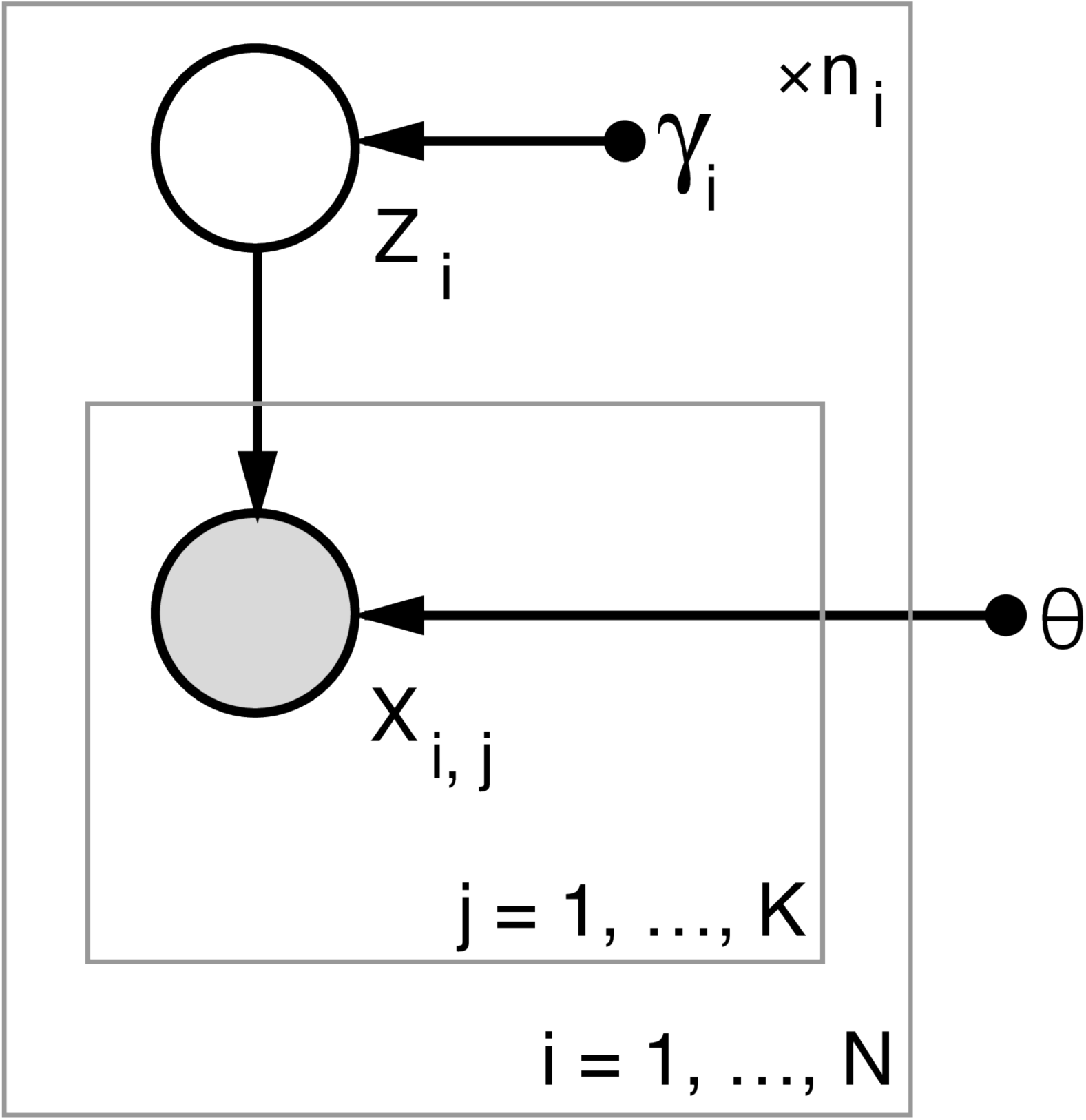
Graphical representation of k-mer mask optimization via enzyme cut-site model. We model the enzyme cut bias as a PSWM, of length K and parameterized by *θ*, shared among across all N possible k-mers. Here *X*_*i,j*_ represents the observed j-th base of the i-th k-mer sequence. Each k-mer has an unknown orientation, represented by the random variable *Z*_*i*_ and parameterized by *γ*_*i*_. Furthermore, each k-mer is observed *n*_*i*_ times in the data. Thus, the full likelihood of the model is:

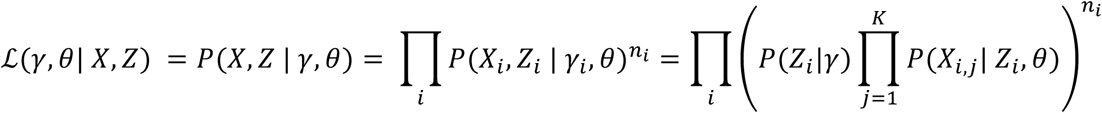

**Figure S2.**
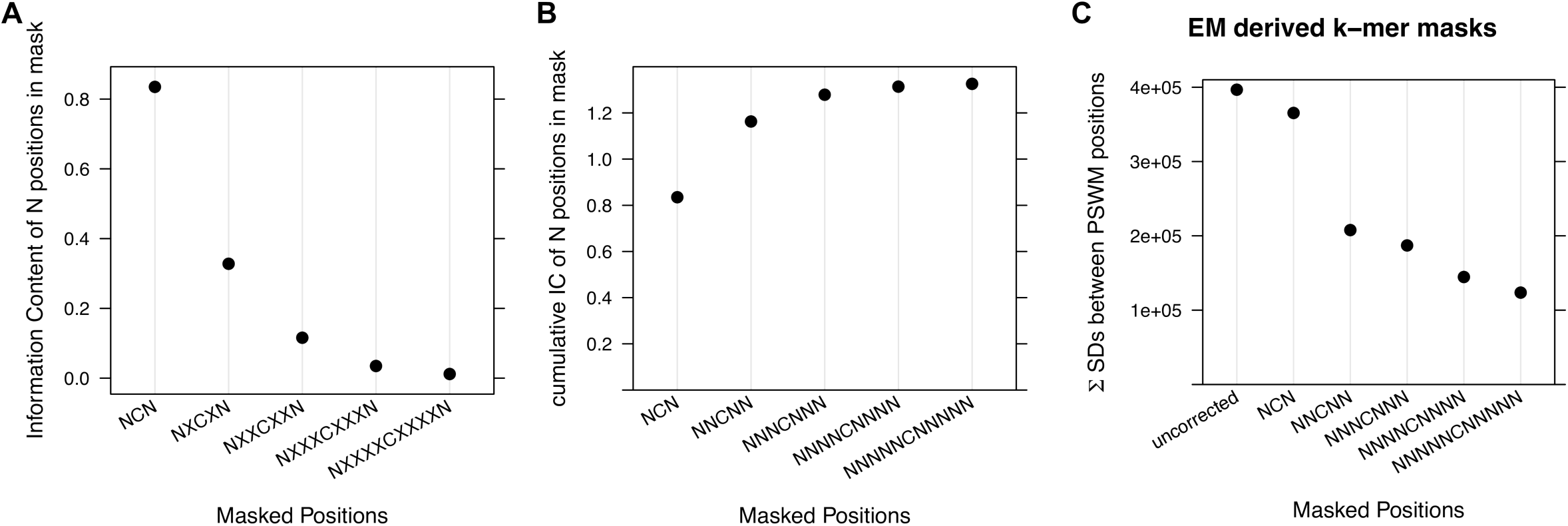
DNase k-mer mask optimization. (A) The PSWM resulting from the expectation maximization k-mer optimization method shows that the information content of the positions gradually decreases with increasing distance from the DNase nick site. (B) The plot of cumulative information content of the positions begins to level off at the tetramer. (C) We plot the decrease in the summed standard deviations across a set of PSWMs as we successively increase the k-mer size relative to the centered DNase nick site.

**Figure S3.**
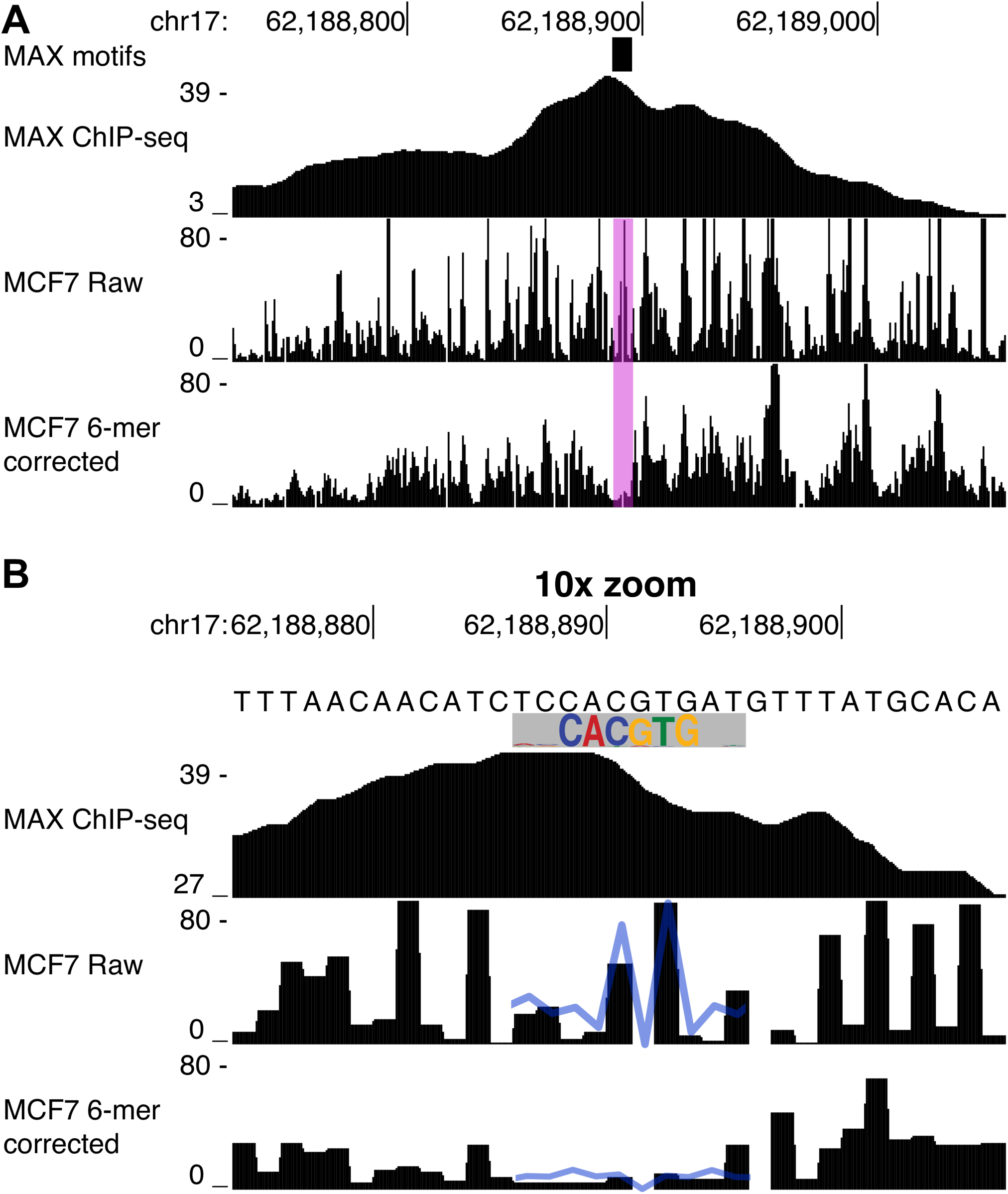
Corrected DNase-seq data reveals footprints at a MAX binding site. (A) The precise location of MAX binding (transparent pink) is inferred from the presence of a MAX recognition motif within a MAX ChIP-seq peak. This binding site shows sharp peaks within the binding site in uncorrected DNase-seq profiles; a footprint is present only in the corrected data. (B) A zoom in of the panel (A) reveals that the composite MAX trace from Figure 4A (shown here as a blue trace) is observed at the MAX binding sites.

**Figure S4.**
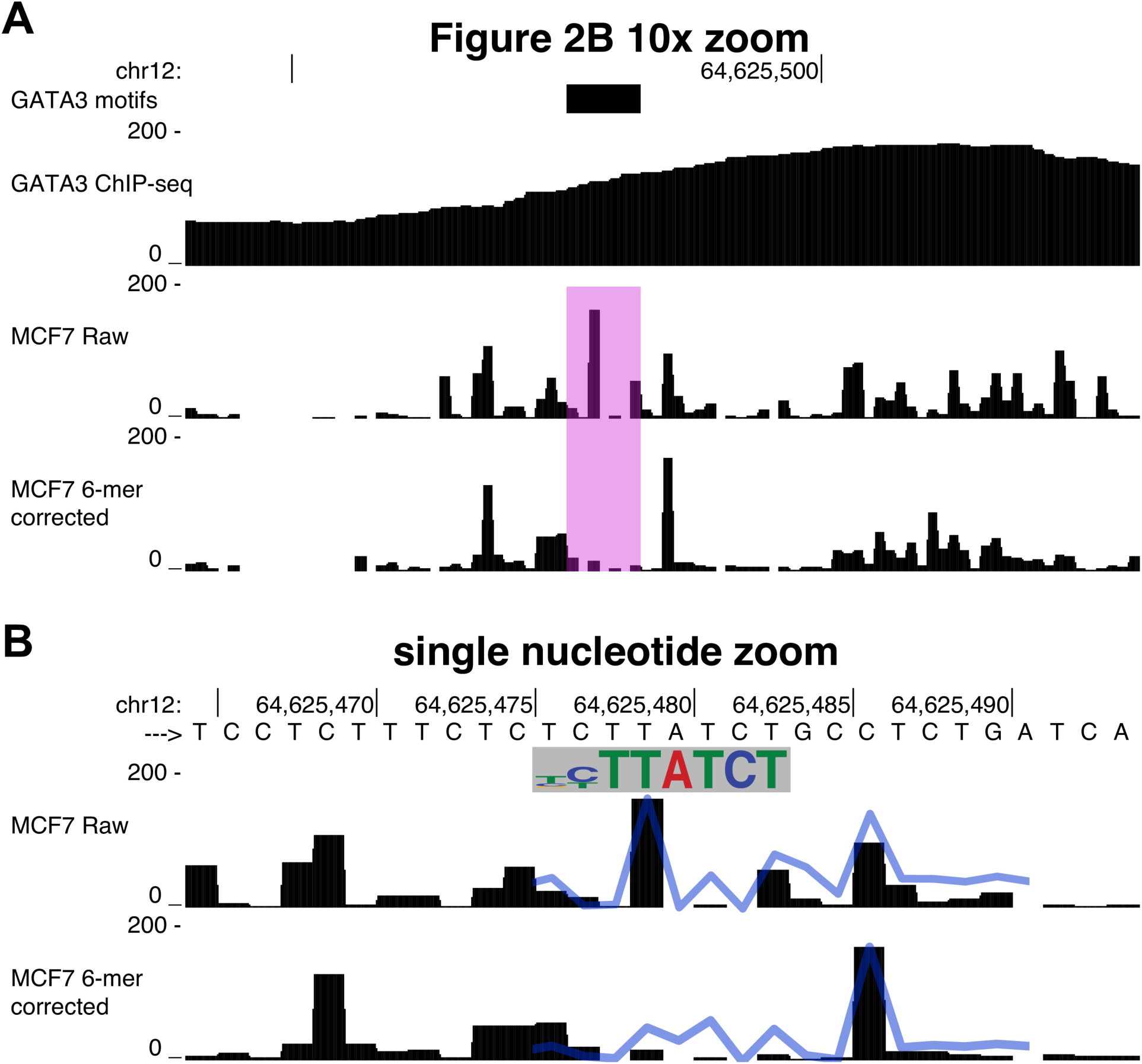
Corrected DNase-seq data reveals footprints at a GATA3 binding site. (A) A 10x zoom in of Figure 2B. (B) The single nucleotide resolution profile reveals that the composite GATA3 trace from Figure 4B (shown here as a blue trace) is observed at the GATA3 binding sites. The molecular signature upstream of GATA3 binding in Figure 4B is observed and enhanced at this binding site at position chr12:64625486.

**Figure S5.**
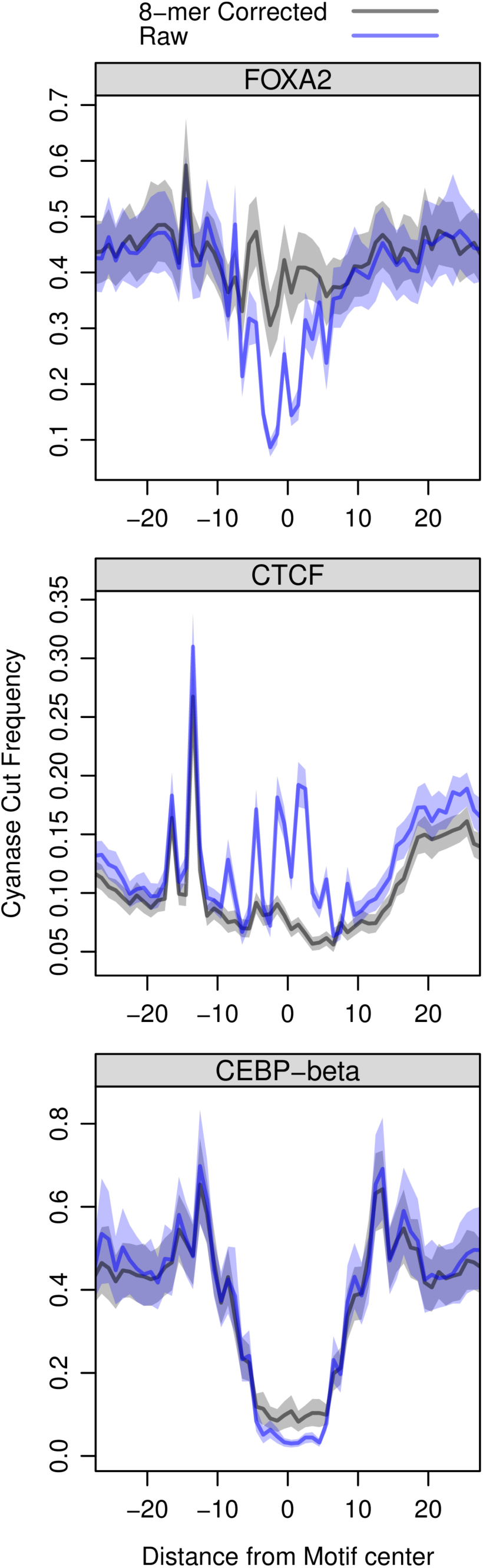
*SeqOutBias* corrects cyanase endonuclease bias. Each composite profile illustrates the average cut frequency at each position between nucleotides. The blue trace is the raw data and the black trace is the 8-mer corrected data.

**Figure S6.**
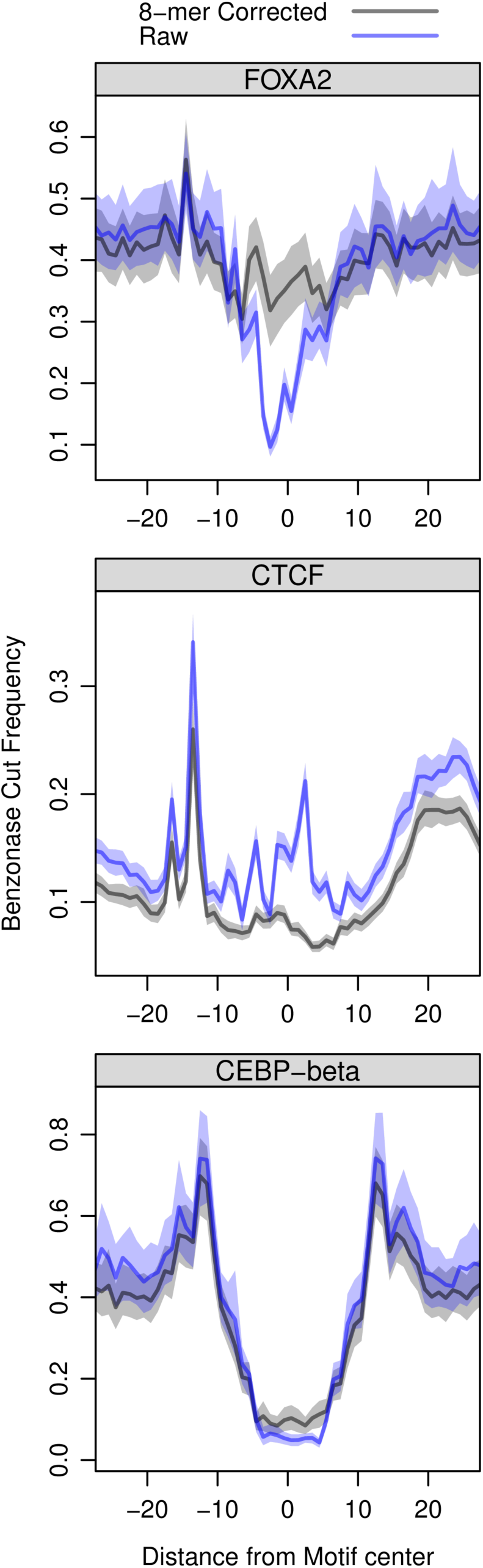
*SeqOutBias* corrects benzonase endonuclease bias. The composite profiles for FOXA2, CTCF, and CEBP-beta binding sites illustrate the average cut frequency at each position between nucleotides. The blue trace is the raw data and the black trace is the 8-mer corrected data.

**Figure S7.**
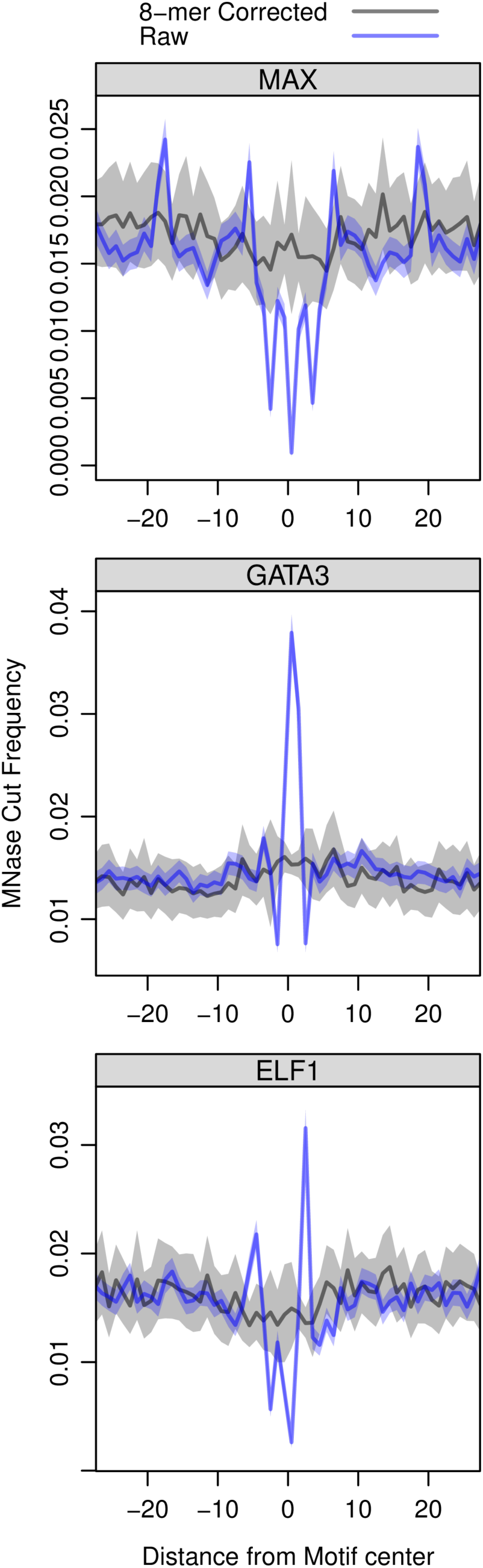
*SeqOutBias* corrects MNase sequence bias. The composite profiles for MAX, GATA3, and ELF1 indicate that sequence correction abrogates the sharp peaks in the traces. The blue trace is the raw data and the black trace is the 8-mer corrected data.

**Figure S8.**
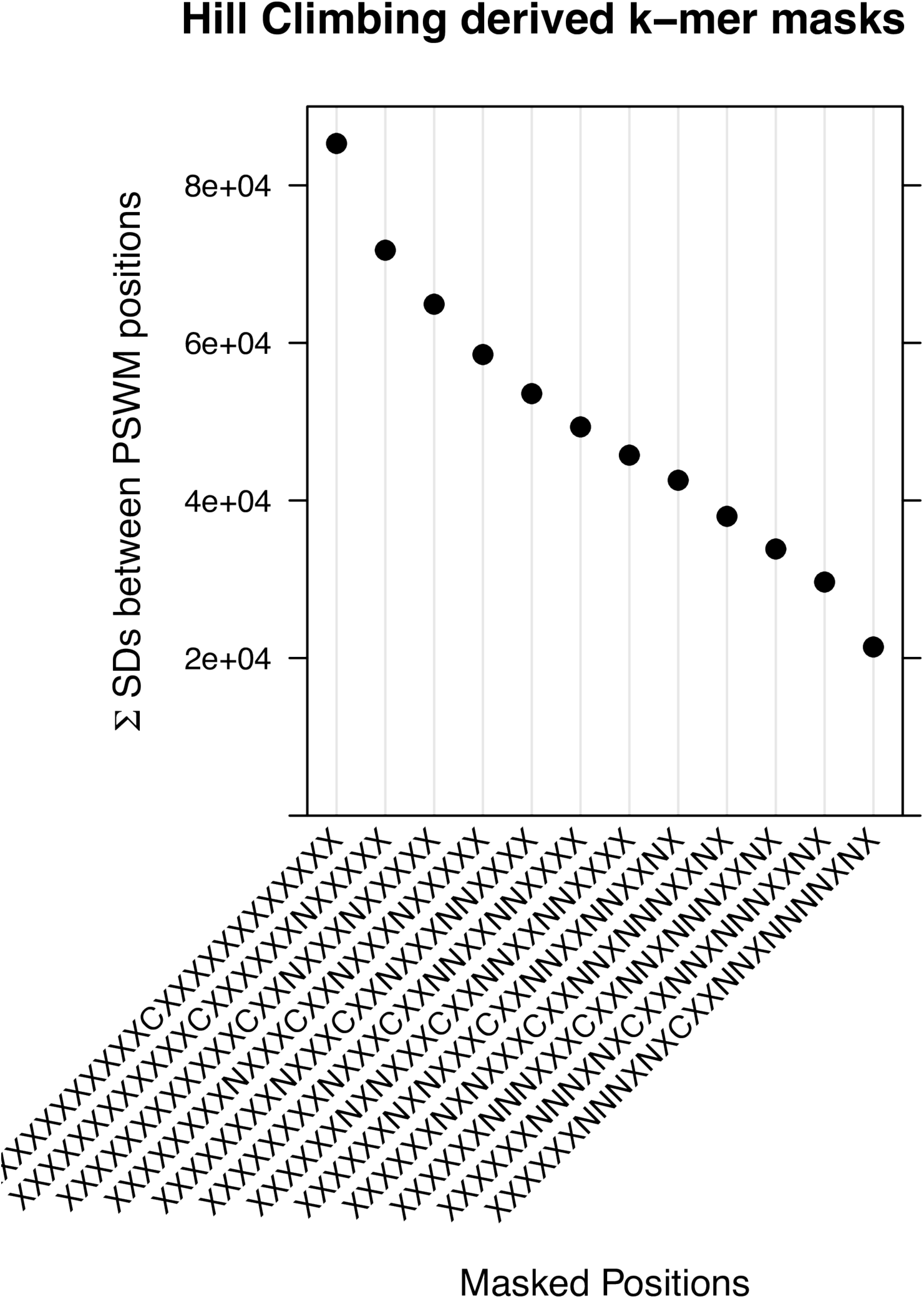
ATAC-seq k-mer mask optimization. We started with a k-mer mask of 12 X bases flanking each side of the Tn5 insertion site and we systematically changed each X position into a masked N. We plot the decrease in the summed standard deviations across a set of PSWMs for the top 11 positions that contribute to Tn5 sequence bias.

**Figure S9.**
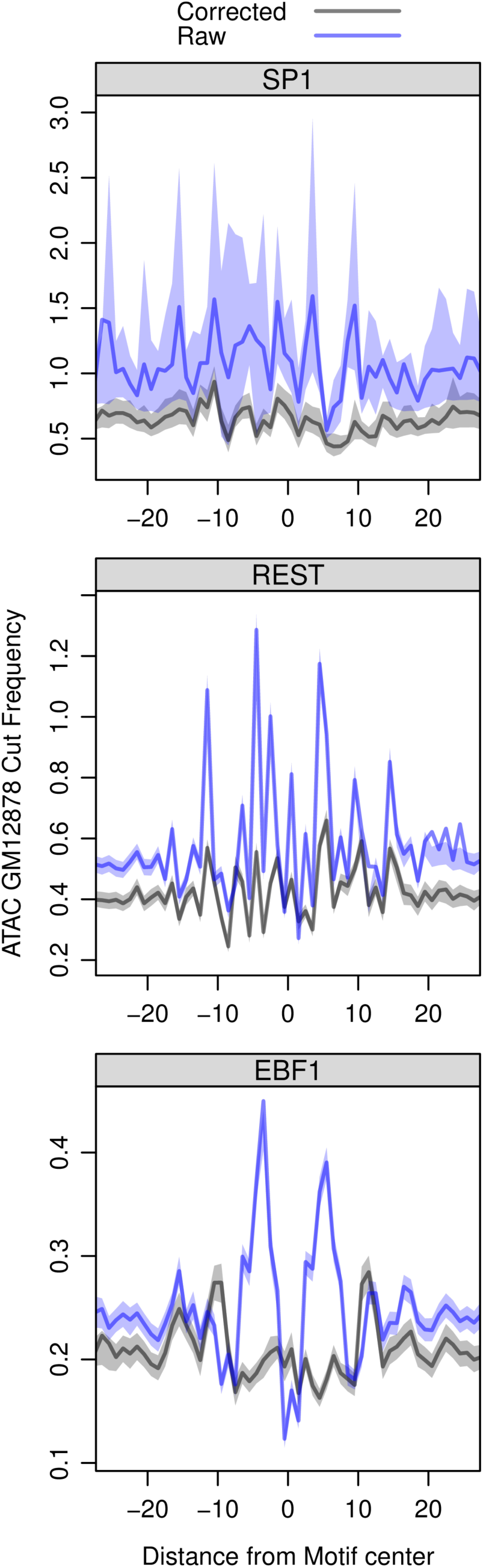
Tn5 insertion bias is corrected in a ATAC-seq experiment from GM12878 cells. The composite profiles for SP1, EBF1, and REST indicate that sequence correction dampens the sharp peaks in the traces.

**Figure S10.**
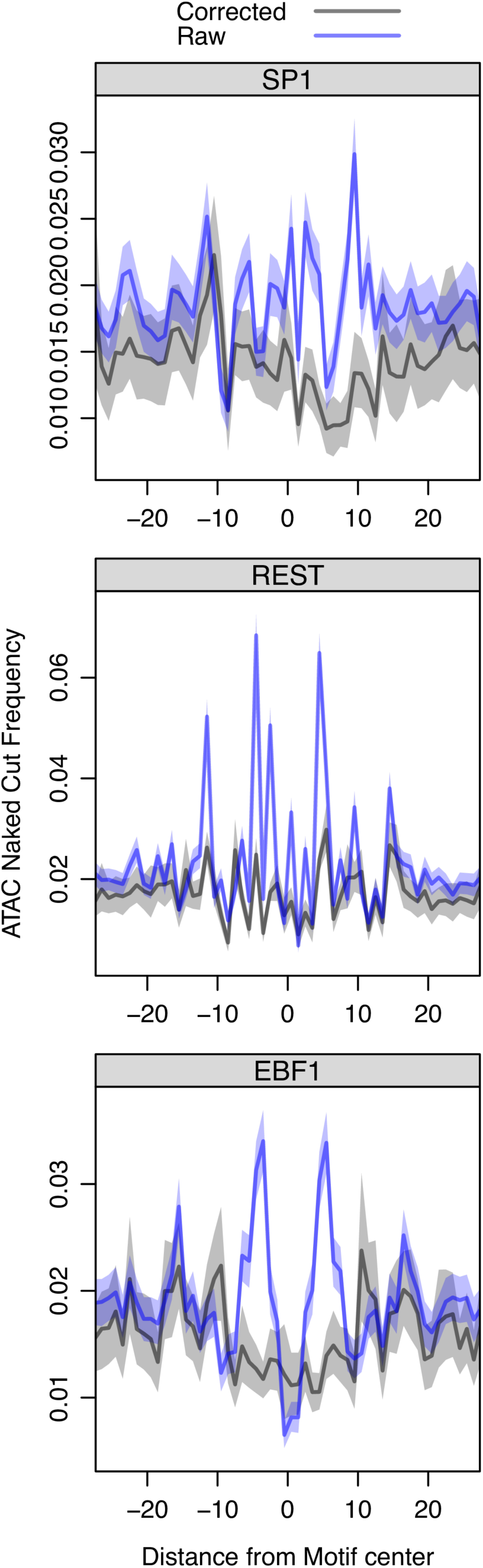
Tn5 insertion bias is corrected in a ATAC-seq experiment from naked DNA. We generated ATAC-seq data with naked DNA and we find that the composite profiles for TFs exhibit dampened sharp peaks in the corrected traces.

**Figure S11.**
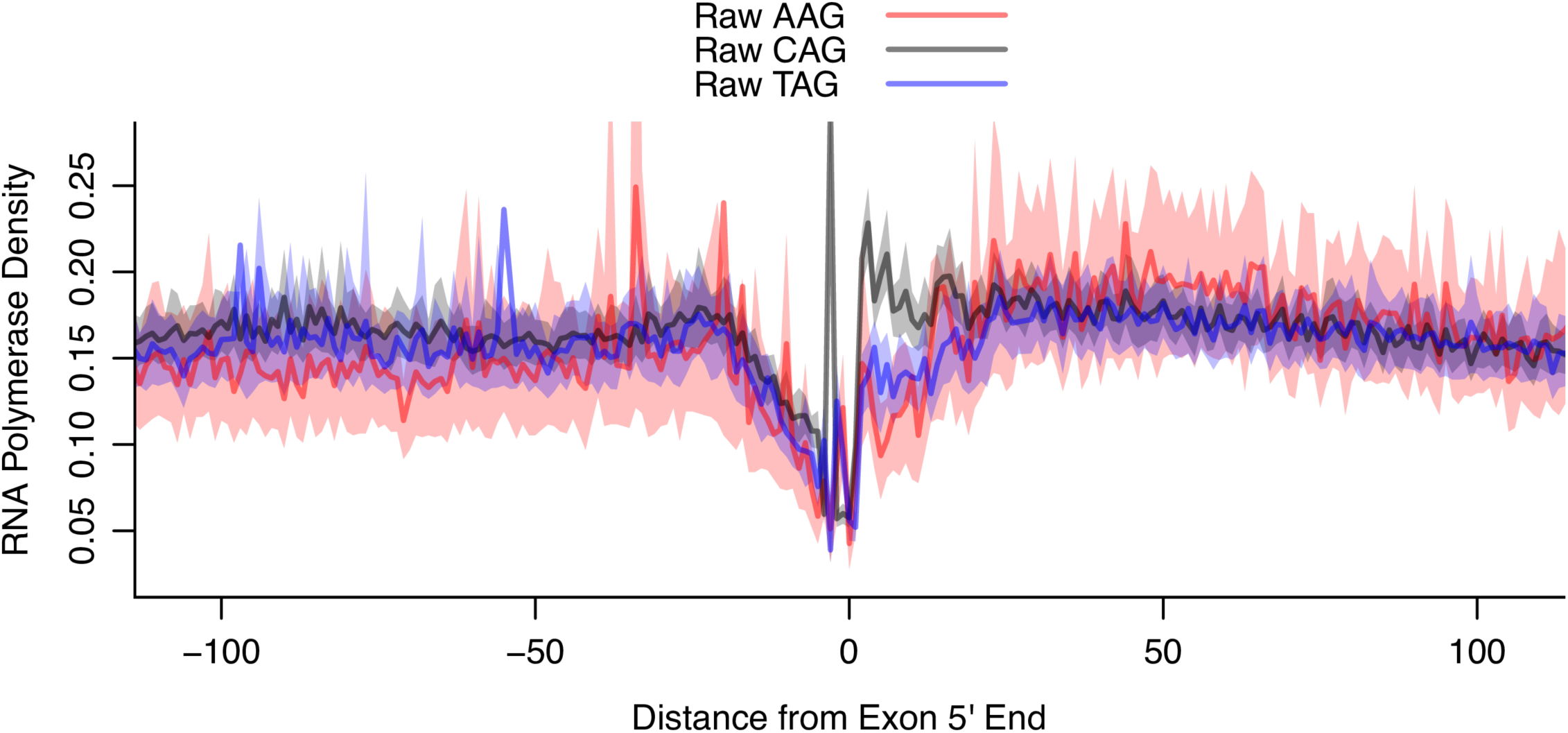
Sequence bias drives a sharp peak of PRO-seq signal upstream of exons. The sharp peak at position −3 found in the CAG splice acceptor profile indicates that cytosine is preferentially incorporated during the nuclear run-on or preferentially ligated during the library preparation.

**Figure S12.**
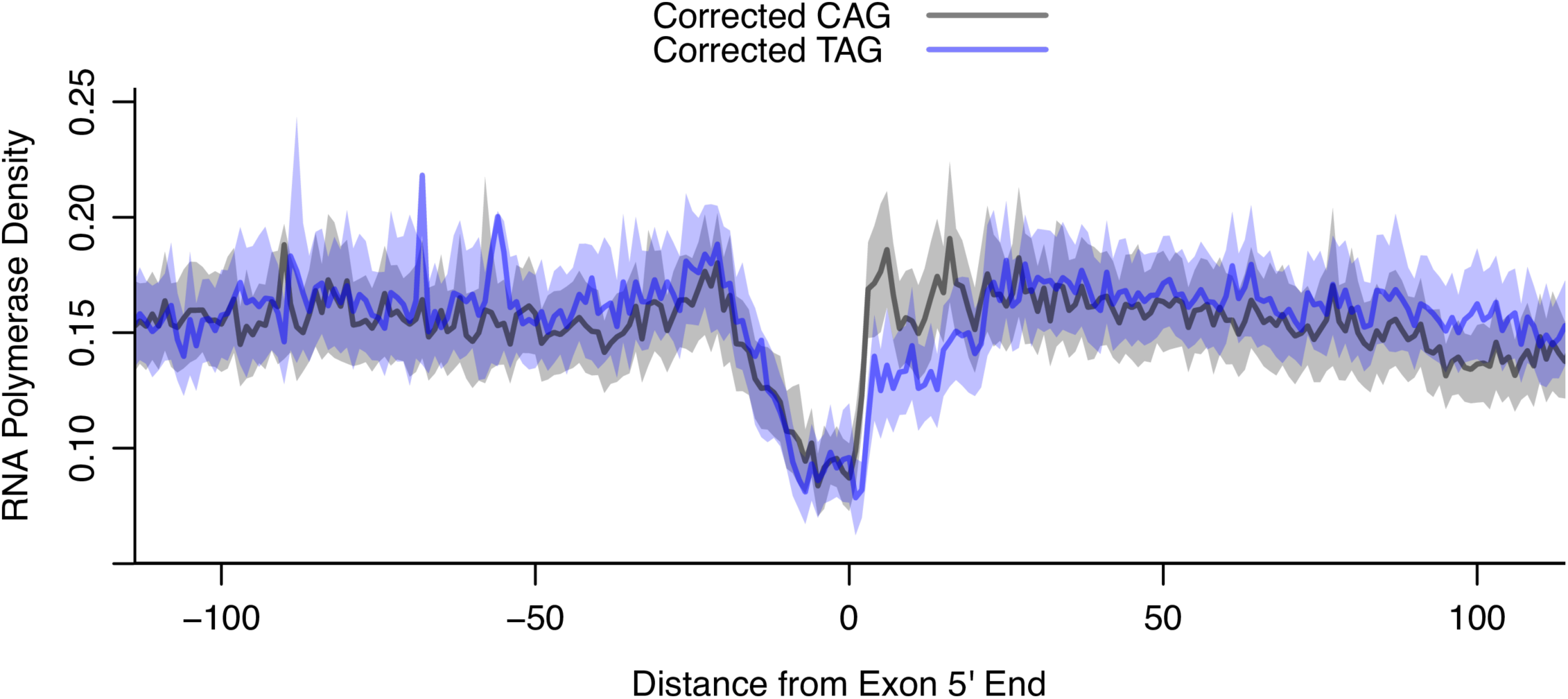
*SeqOutBias* corrects PRO-seq sequence bias. Corrected PRO-seq profiles abrogate the sharp peak at position −3 at CAG consensus exons and the modest difference in intensity in the 5´end of the exon indicates that RNA Polymerase may proceed faster at TAG exons relative to CAG.

**Figure S13.**
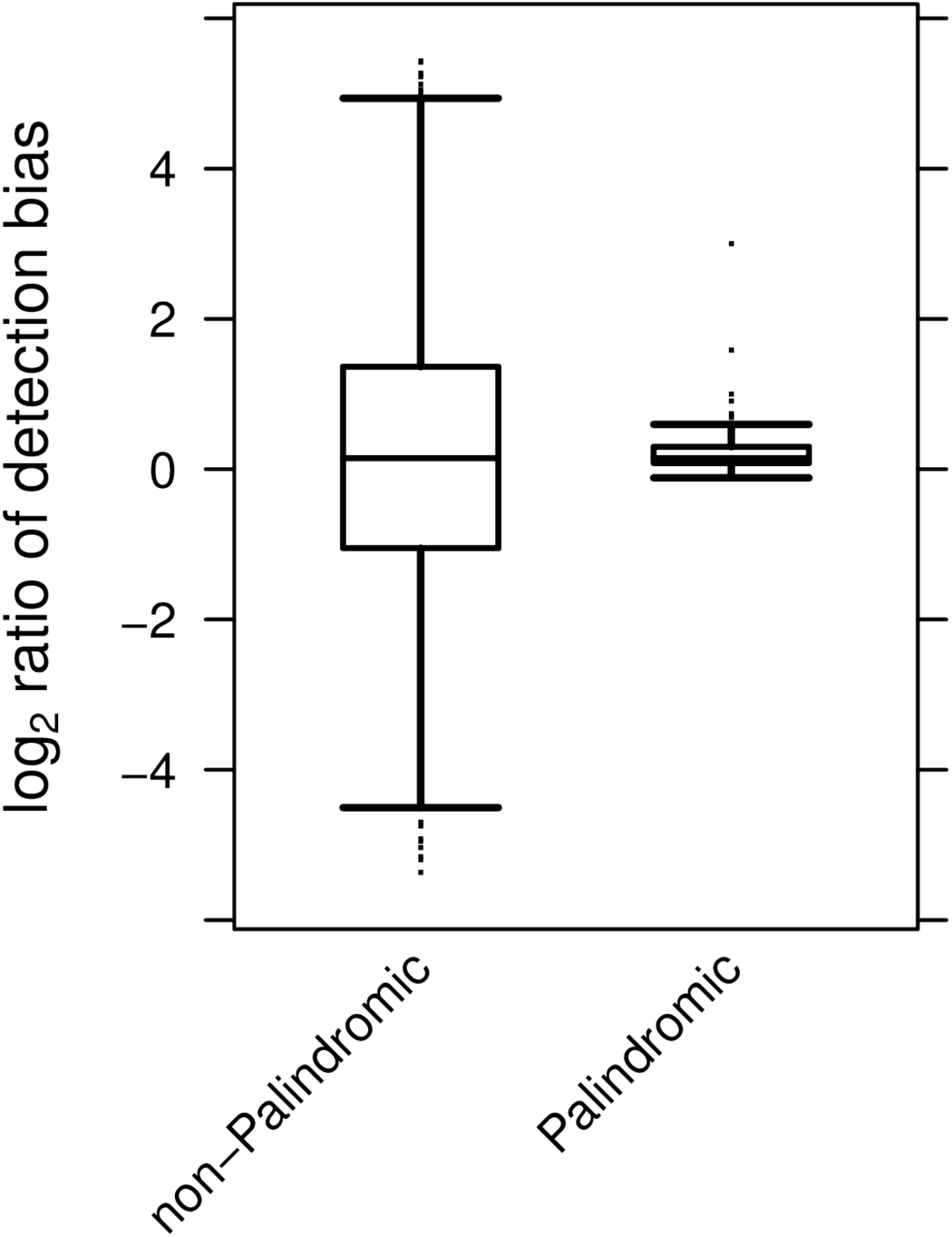
Reverse palindromic hexamers do not exhibit enzymatic DNA end repair and ligation bias. The DNA end substrates are identical in reverse palindromic nick sites, accounting for the absence of bias.

**Figure S14.**
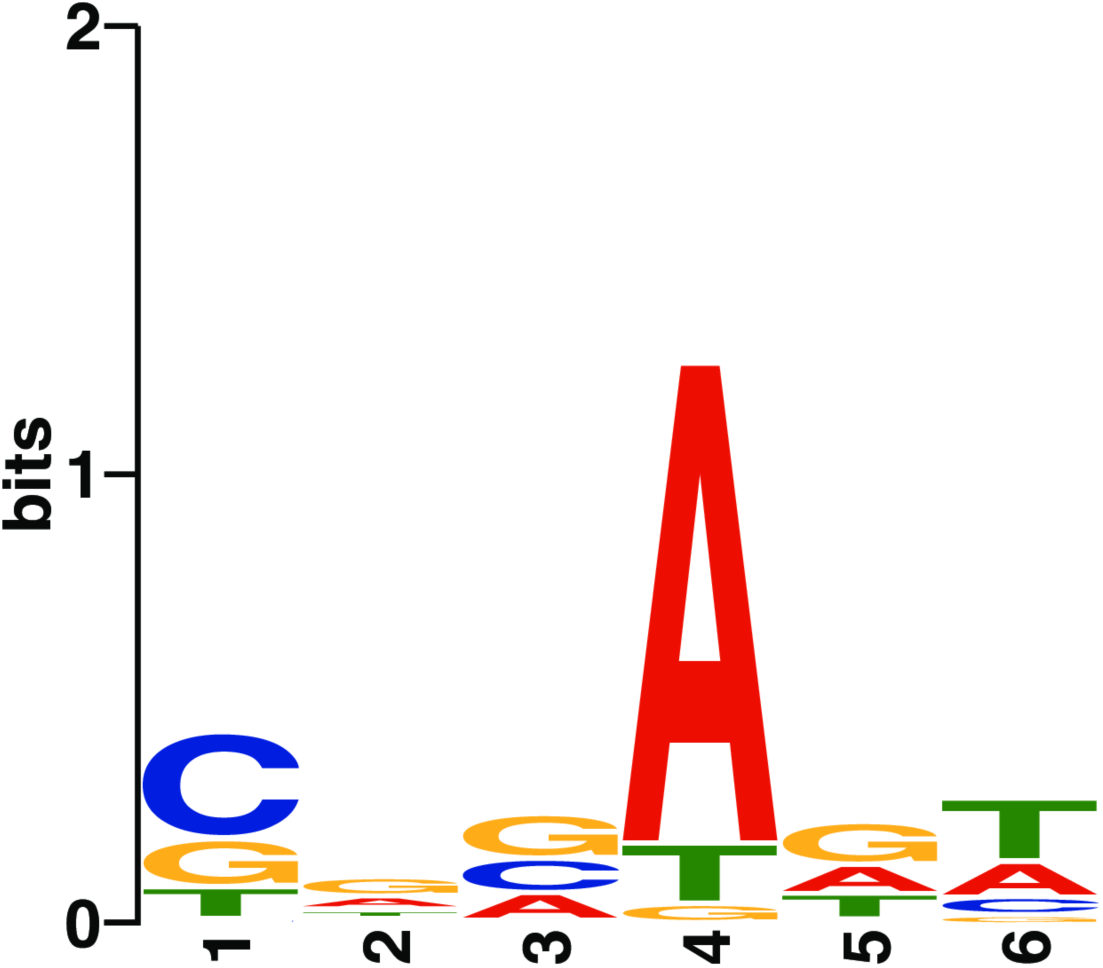
DNA sequence drives DNA end repair and ligation preference. Motif analysis of the 5% most enzymatic DNA end repair and ligation biased 6-mers indicates that an Adenine in position 4 of the 6-mer is preferentially sequenced compared to the oppositely oriented nucleotide in position 3.

**Figure S15.**
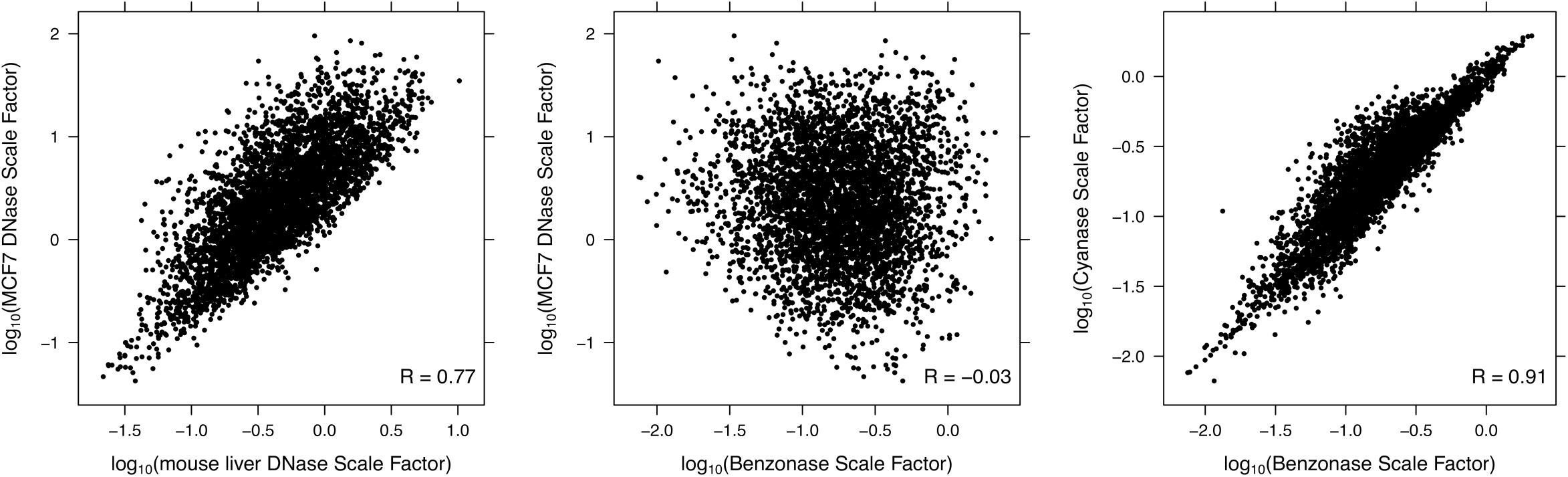
Enzymatic nick biases are correlated between DNase-seq experiments and correlated between Cyanase and Benzonase digestion experiments. These scatter plots show that the enzymatic nick biases, as measured by the *seqOutBias* scale factor, are correlated between DNase experiments and correlated between Cyanase and Benzonase.

**Figure S16.**
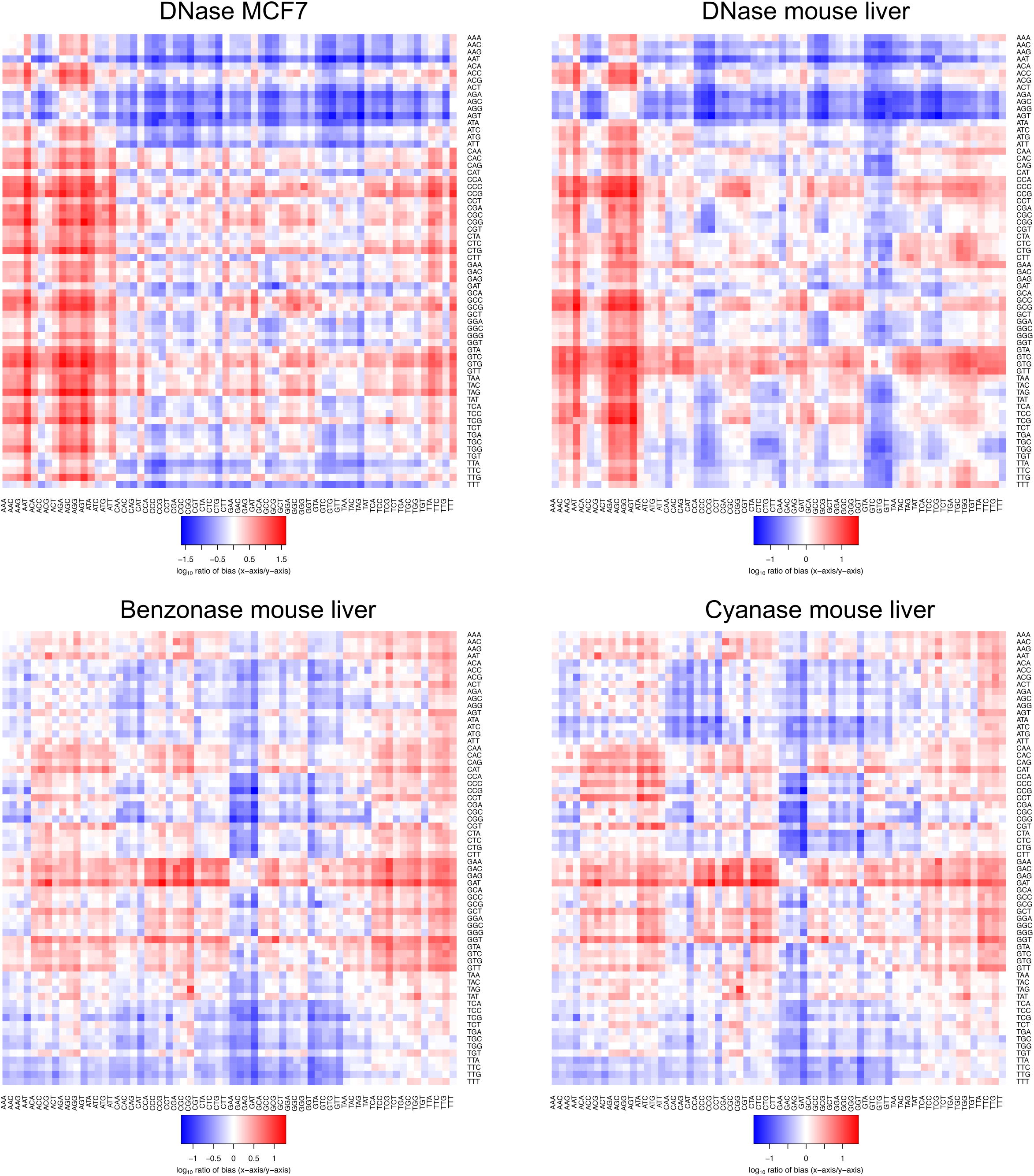
Post-nick enzymatic processing biases of DNase are correlated between experiments and the post nick biases of Cyanase and Benzonase are similar. The relative bias of all 3-mers sequenced (the ratio of x-axis 3-mer to y-axis 3-mer) for four separate experiments.

## References

1. Hesselberth,J.R., Chen,X., Zhang,Z., Sabo,P.J., Sandstrom,R., Reynolds,A.P., Thurman,R.E., Neph,S., Kuehn,M.S., Noble,W.S., et al. (2009) Global mapping of protein-DNA interactions in vivo by digital genomic footprinting. Nat. Methods, 6, 283–289.

2. Boyle,A.P., Davis,S., Shulha,H.P., Meltzer,P., Margulies,E.H., Weng,Z., Furey,T.S. and Crawford,G.E. (2008) High-resolution mapping and characterization of open chromatin across the genome. Cell, 132, 311–322.

3. Rhee,H.S. and Pugh,B.F. (2011) Comprehensive genome-wide protein-DNA interactions detected at single-nucleotide resolution. Cell, 147, 1408–1419.

4. Kwak,H., Fuda,N.J., Core,L.J. and Lis,J.T. (2013) Precise maps of RNA polymerase reveal how promoters direct initiation and pausing. Science, 339, 950–953.

5. Duarte,F.M., Fuda,N.J., Mahat,D.B., Core,L.J., Guertin,M.J. and Lis,J.T. (2016) Transcription factors GAF and HSF act at distinct regulatory steps to modulate stress-induced gene activation. Genes Dev., 30, 1731–1746.

6. Grøntved,L., Bandle,R., John,S., Baek,S., Chung,H.-J., Liu,Y., Aguilera,G., Oberholtzer,C., Hager,G.L. and Levens,D. (2012) Rapid genome-scale mapping of chromatin accessibility in tissue. Epigenetics Chromatin, 5, 10.

7. Buenrostro,J.D., Giresi,P.G., Zaba,L.C., Chang,H.Y. and Greenleaf,W.J. (2013) Transposition of native chromatin for fast and sensitive epigenomic profiling of open chromatin, DNA-binding proteins and nucleosome position. Nat. Methods, 10, 1213–1218.

8. Wu,C., Bingham,P.M., Livak,K.J., Holmgren,R. and Elgin,S.C. (1979) The chromatin structure of specific genes: I. Evidence for higher order domains of defined DNA sequence. Cell, 16, 797–806.

9. Neph,S., Vierstra,J., Stergachis,A.B., Reynolds,A.P., Haugen,E., Vernot,B., Thurman,R.E., John,S., Sandstrom,R., Johnson,A.K., et al. (2012) An expansive human regulatory lexicon encoded in transcription factor footprints. Nature, 489, 83–90.

10. Sung,M.-H., Guertin,M.J., Baek,S. and Hager,G.L. (2014) DNase footprint signatures are dictated by factor dynamics and DNA sequence. Mol. Cell, 56, 275–285.

11. He,H.H., Meyer,C.A., Hu,S.S., Chen,M.-W., Zang,C., Liu,Y., Rao,P.K., Fei,T., Xu,H., Long,H., et al. (2014) Refined DNase-seq protocol and data analysis reveals intrinsic bias in transcription factor footprint identification. Nat. Methods, 11, 73–78.

12. Yardimci,G.G., Frank,C.L., Crawford,G.E. and Ohler, U. (2014) Explicit DNase sequence bias modeling enables high-resolution transcription factor footprint detection. Nucleic Acids Res., 42, 11865–11878.

13. Piper,J., Elze,M.C., Cauchy,P., Cockerill,P.N., Bonifer,C. and Ott,S. (2013) Wellington: a novel method for the accurate identification of digital genomic footprints from DNase-seq data. Nucleic Acids Res., 41, e201.

14. Gusmao,E.G., Allhoff,M., Zenke,M. and Costa,I.G. (2016) Analysis of computational footprinting methods for DNase sequencing experiments. Nat. Methods, 13, 303–309.

15. Piper,J., Assi,S.A., Cauchy,P., Ladroue,C., Cockerill,P.N., Bonifer,C. and Ott,S. (2015) Wellington-bootstrap: differential DNase-seq footprinting identifies cell-type determining transcription factors. BMC Genomics, 16, 1000.

16. Stergachis,A.B., Neph,S., Reynolds,A., Humbert,R., Miller,B., Paige,S.L., Vernot,B., Cheng,J.B., Thurman,R.E., Sandstrom,R., et al. (2013) Developmental fate and cellular maturity encoded in human regulatory DNA landscapes. Cell, 154, 888–903.

17. Gremme,G., Steinbiss,S. and Kurtz,S. (2013) GenomeTools: a comprehensive software library for efficient processing of structured genome annotations. IEEE/ACM Trans. Comput. Biol. Bioinform., 10, 645–656.

18. Kurtz,S., Narechania,A., Stein,J.C. and Ware,D. (2008) A new method to compute K-mer frequencies and its application to annotate large repetitive plant genomes. BMC Genomics, 9, 517.

19. Thurman,R.E., Rynes,E., Humbert,R., Vierstra,J., Maurano,M.T., Haugen,E., Sheffield,N.C., Stergachis,A.B., Wang,H., Vernot,B., et al. (2012) The accessible chromatin landscape of the human genome. Nature, 489, 75–82.

20. Wu,C., Wong,Y.C. and Elgin,S.C. (1979) The chromatin structure of specific genes: II. Disruption of chromatin structure during gene activity. Cell, 16, 807–814.

21. Wu,C. (1980) The 5' ends of Drosophila heat shock genes in chromatin are hypersensitive to DNase I. Nature, 286, 854–860.

22. Kähärä,J. and Lähdesmäki,H. (2015) BinDNase: a discriminatory approach for transcription factor binding prediction using DNase I hypersensitivity data. Bioinformatics, 31, 2852–2859.

23. Lazarovici,A., Zhou,T., Shafer,A., Dantas Machado,A.C., Riley,T.R., Sandstrom,R., Sabo,P.J., Lu,Y., Rohs,R., Stamatoyannopoulos,J.A., et al. (2013) Probing DNA shape and methylation state on a genomic scale with DNase I. Proc. Natl. Acad. Sci. U. S. A., 110, 6376–6381.

24. Boyle,A.P., Song,L., Lee,B.-K., London,D., Keefe,D., Birney,E., Iyer,V.R., Crawford,G.E. and Furey,T.S. (2011) High-resolution genome-wide in vivo footprinting of diverse transcription factors in human cells. Genome Res., 21, 456–464.

25. Grøntved,L., John,S., Baek,S., Liu,Y., Buckley,J.R., Vinson,C., Aguilera,G. and Hager,G.L. (2013) C/EBP maintains chromatin accessibility in liver and facilitates glucocorticoid receptor recruitment to steroid response elements. EMBO J., 32, 1568–1583.

26. Soccio, R.E., Tuteja,G., Everett,L.J., Li,Z., Lazar,M.A. and Kaestner,K.H. (2011) Species-specific strategies underlying conserved functions of metabolic transcription factors. Mol. Endocrinol., 25, 694–706.

27. Stamatoyannopoulos,J.A., Snyder,M., Hardison,R., Ren,B., Gingeras,T., Gilbert,D.M., Groudine,M., Bender,M., Kaul,R., Canfield,T., et al. (2012) An encyclopedia of mouse DNA elements (Mouse ENCODE). Genome Biol., 13, 418.

28. Guertin,M.J., Zhang,X., Coonrod,S.A. and Hager,G.L. (2014) Transient estrogen receptor binding and p300 redistribution support a squelching mechanism for estradiol-repressed genes. Mol. Endocrinol., 28, 1522– 1533.

29. ENCODE Project Consortium (2012) An integrated encyclopedia of DNA elements in the human genome. Nature, 489, 57–74.

30. Core,L.J., Martins,A.L., Danko,C.G., Waters,C.T., Siepel,A. and Lis,J.T. (2014) Analysis of nascent RNA identifies a unified architecture of initiation regions at mammalian promoters and enhancers. Nat. Genet., 46, 1311–1320.

31. Core,L.J., Waterfall,J.J. and Lis,J.T. (2008) Nascent RNA sequencing reveals widespread pausing and divergent initiation at human promoters. Science, 322, 1845–1848.

32. Sickmier,E.A., Frato,K.E., Shen,H., Paranawithana,S.R., Green,M.R. and Kielkopf,C.L. (2006) Structural basis for polypyrimidine tract recognition by the essential pre-mRNA splicing factor U2AF65. Mol. Cell, 23, 49–59.

33. Guth,S. and Valcárcel,J. (2000) Kinetic role for mammalian SF1/BBP in spliceosome assembly and function after polypyrimidine tract recognition by U2AF. J. Biol. Chem., 275, 38059–38066.

34. Homan,P.J., Tandon,A., Rice,G.M., Ding,F., Dokholyan,N.V. and Weeks,K.M. (2014) RNA tertiary structure analysis by 2’-hydroxyl molecular interference. Biochemistry, 53, 6825–6833.

